# Enhancer cooperativity can compensate for loss of activity over large genomic distances

**DOI:** 10.1101/2023.12.06.570399

**Authors:** Henry Thomas, Songjie Feng, Marie Huber, Vincent Loubiere, Daria Vanina, Mattia Pitasi, Alexander Stark, Christa Buecker

## Abstract

Enhancers are short DNA sequences that activate their target promoter from a distance; however, increasing the genomic distance between the enhancer and the promoter decreases expression levels. Many genes are controlled by combinations of multiple enhancers, yet the interaction and cooperation of individual enhancer elements is not well understood. Here, we developed a novel synthetic platform that allows building complex regulatory landscapes from the bottom up. We tested the system by integrating individual enhancers at different distances and revealed that the strength of an enhancer determines how strongly it is affected by increased genomic distance. Furthermore, synergy between two enhancer elements depends on the distance at which the two elements are integrated: introducing a weak enhancer between a strong enhancer and the promoter strongly increases reporter gene expression, allowing enhancers to activate from increased genomic distances.

## Introduction

The transcription of mRNA molecules from the promoter of a gene is the first step in gene expression that needs to be tightly controlled to ensure correct expression levels. In higher eukaryotes, cis-regulatory elements such as enhancers control when and in which cells a promoter is active (Kim and Wysocka 2023; Long, Prescott, and Wysocka 2016). Enhancers are short DNA sequences that consist of multiple transcription factor binding sites and control the expression of their target genes from a distance (Long, Prescott, and Wysocka 2016). However, with increasing genomic distance, the activation potential of an individual enhancer on the promoter decreases (Zuin et al. 2022; Rinzema et al. 2022), raising the question of how enhancers can bridge the distance to their promoter.

Enhancers are typically identified and studied either in their native genomic context or in reporter assays (Long, Prescott, and Wysocka 2016; Catarino and Stark 2018). Both approaches have contributed remarkably to our understanding of how enhancers activate target gene expression. Detailed dissection of enhancer loci has e. g. demonstrated the importance of tissue-specific TFs (Kim and Wysocka 2023; Buecker et al. 2014), identified a role for 3D genome organisation in facilitating enhancer-promoter communication (Long, Prescott, and Wysocka 2016; Pachano, Haro, and Rada-Iglesias 2022) and described various modes of enhancer cooperativity (Bothma et al. 2015; Hay et al. 2016; Carleton, Berrett, and Gertz 2017; Huang et al. 2016; Shin et al. 2016; Hnisz et al. 2015; Moorthy et al. 2017; Thomas et al. 2021; Brosh et al. 2023). However, the complexity of endogenous regulatory landscapes can obstruct their analysis. A recent dissection of the ⍺-globin locus used advances in synthesis, manipulation, and integration of entire genomic loci to remove all enhancers at the locus and re-introduce them one by one (Blayney et al. 2022). Importantly, this approach revealed the super-additive behaviour of several enhancer elements, contrasting previous conclusions of additive behaviour at the same locus based on deleting individual enhancer elements (Hay et al. 2016; Blayney et al. 2022). This outcome underlines the importance of studying individual elements of multi-enhancer clusters both in the presence and absence of other cis-regulatory elements to understand their function and their cooperative behaviour fully. Genetic modification of endogenous enhancer landscapes remains laborious, and using such an approach is limited to a few selected enhancer clusters and the constituent of those clusters, making it challenging to derive general rules about how different enhancers can cooperate.

Conversely, reporter assays can eliminate regulatory complexity and quickly analyse the activity of individual enhancers and even combinations. Besides testing in which tissue a given enhancer is active and how strongly it activates transcription (Catarino and Stark 2018; Gasperini, Tome, and Shendure 2020), reporter assays can be massively parallelised (massively parallel reporter assays, MPRAs) to measure the enhancer activity of millions of fragments in one experiment and identify genomic fragments with enhancer activity in the cell type of interest at ease (Arnold et al. 2013; Smith et al. 2013). Different classes of enhancers with distinct promoter compatibility or co-factor dependency have been identified by combining enhancers with different promoters or using MPRAs with perturbation of regulatory factors (Neumayr et al. 2022; Zabidi et al. 2015). However, one major limitation of MPRAs is the size of the analysed DNA fragment. Therefore, enhancers are typically placed very close to the promoter, so enhancer-promoter communication over large genomic distances cannot be investigated (Muerdter et al. 2018). Furthermore, most genes are controlled by multiple enhancers that activate the promoter together (Gschwind et al. 2023). While first studies combined two enhancers to study cooperativity in their activation of a single target gene (Loubiere et al. 2023; Martinez-Ara, Comoglio, and Steensel 2023), the genomic distance between the elements was negligible compared to the distances enhancers have to overcome in mammalian genomes. Finally, elements with low or undetectable enhancer activity in reporter assays have been shown to exert critical regulatory functions in combination with other enhancers at their respective endogenous loci (Blayney et al. 2022; Batut et al. 2022; Thomas et al. 2021; L.-F. Chen et al. 2023). Therefore, reporter assays cannot fully recapitulate all aspects of endogenous gene regulation, and new experimental systems are required to enable the study of enhancers in their native environment with higher throughput as compared to previous dissections of enhancer clusters at their endogenous loci.

Here, we developed a novel platform for testing enhancer activities and their distance dependencies: a cell line that allows for efficient integration of enhancer sequences at three different distances to a fluorescent reporter gene. Due to the three integration sites and the fact that this system has been placed in an empty TAD without additional regulatory elements, we can build and analyse complex regulatory landscapes from the bottom up. The relative ease with which enhancers can be integrated into the three integration sites allowed us to compare five enhancers selected from different genomic contexts of the mouse genome at different distances to the same promoter. We demonstrate that all enhancers lose the ability to activate the minimal promoter with increasing distance. However, strong enhancers are less affected by genomic distance. Finally, we show that combinations of weak and strong enhancers strongly increase expression levels when the weak enhancer is placed between the strong enhancer and the promoter, but not when the weak enhancer is placed upstream of the strong enhancer. Thus, synergy between enhancers depends not only on the individual enhancer activity but also on the genomic distance between the enhancers and the promoter.

## Results

### Design of the reporter system

We aimed to generate a flexible and quantitative reporter system that allows analysis of individual cis-regulatory elements and combinations thereof at different genomic distances to a reporter gene. To reduce the complexity of the regulatory landscape, we decided to introduce the system into an empty TAD containing no active cis-regulatory elements in mESCs. This allows us to focus on the effect of the introduced enhancers on reporter gene expression without the need to account for existing endogenous cis-regulatory elements. We chose the β-globin locus that has been described before as an inert environment in mESCs (Lienert et al. 2011) and is devoid of activating or repressive regulatory elements based on the absence of H3K4me1/3, H3K27ac, H3K27me3 or H3K9me2 histone marks in ESCs (Lienert et al. 2011; Yang et al. 2019). To facilitate enhancer integration and eliminate the need to screen for homozygous integration events, we introduced the reporter locus on only one of the two β-globin alleles. Thus, we used v6.5 mouse ESCs (Rideout et al. 2000) that were derived from a C57BL/6 × 129/sv cross and contained single nucleotide polymorphisms (SNPs) distinguishing the two parental alleles to manipulate only the C57BL/6 allele (Figure 1A). We introduced an mCherry reporter gene under the control of a minimal TK-promoter (Greenshpan et al. 2022; Arcot, Flemington, and Deininger 1989) into the β-globin locus 1.5kb upstream of the Hbb-γ promoter. In addition, we integrated three landing pads (LPs) 1.5 kb, 25 kb and 75 kb upstream of the reporter gene (Fig. 1A). After the introduction of the first reporter, we chose two independent clones that we used for the integration of the additional LPs. All initial enhancer manipulations (Figures 1 and 2) were carried out in both cell lines, reaching the same conclusions.

**Figure 1:**
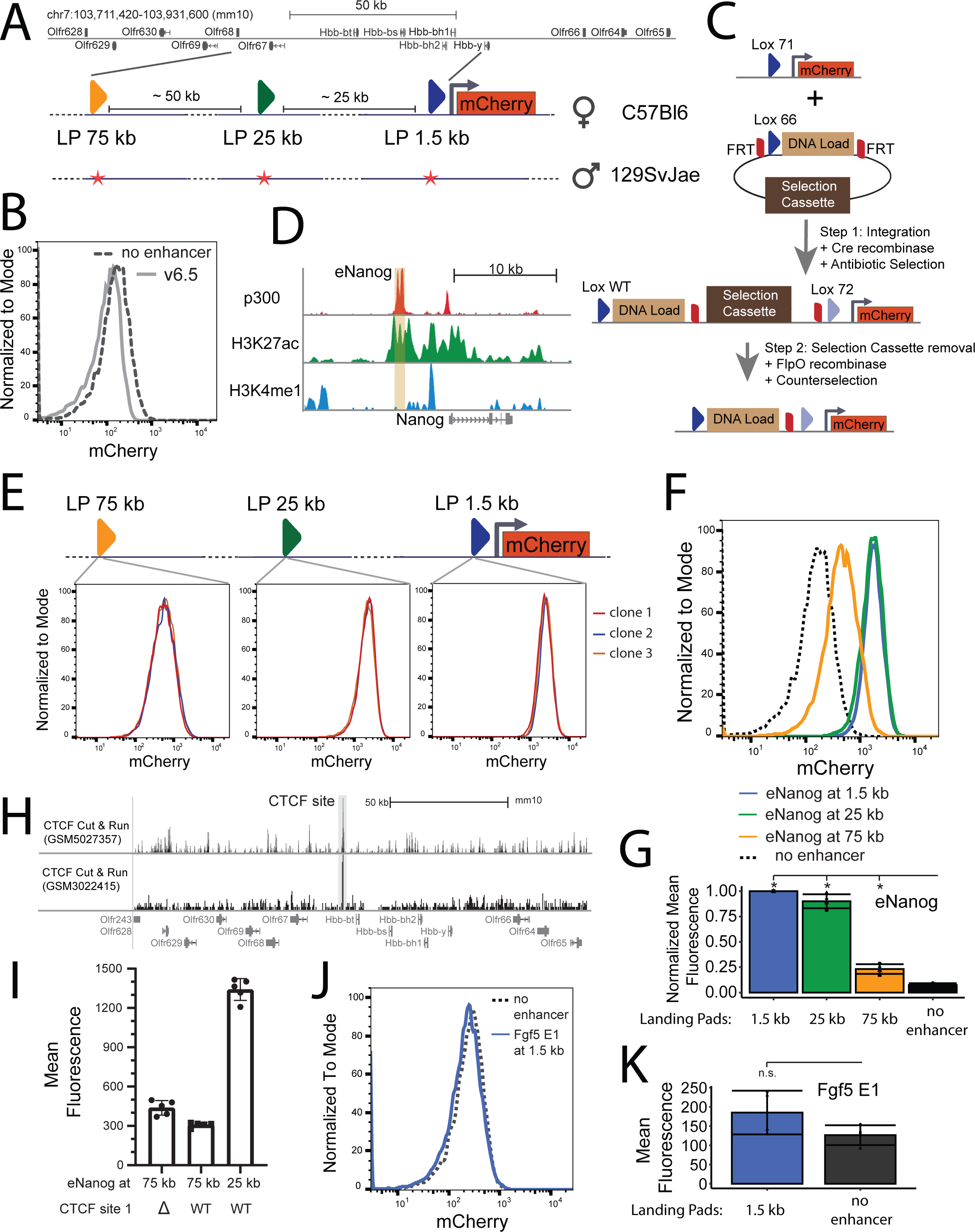
Generation and Validation of a synthetic locus to test enhancer activity systematically. A: Schematic representation of the different components of the synthetic locus generated in this study. Top: UCSC browser track depicting the β-globin locus. Below: different parts of the synthetic locus including three landing pads (LP) at different distances to the minimal promoter (arrow) and the mCherry reporter gene. All components are integrated on the C57Bl6 allele. Bottom: Red stars depict single nucleotide polymorphisms (SNPs) used to target the C57Bl6 allele selectively. B: FACS plot showing a selected example of clone 1 no enhancer and the parental cell line v6.5 C: Schematic representation of the integration strategy (see text for details) D: UCSC browser track depicting p300, H3K27ac, and H3K4me1 ChIP-seq tracks for the *Nanog* locus (Buecker et al. 2014) E: Representative FACS plots showing three individual clones with the Nanog enhancer (eNanog) integrated at the three LPs. F: FACS plots of individual examples for mCherry expression of *eNanog* integrated at each distance. G: Quantification of mean mCherry expression of *eNanog* integrated at the indicated landing pads. All mean expressions were normalised to the 1.5 kb integration. Statistically significant different signals (p < 0.05, one-sided paired t-tests) are marked by stars. H: UCSC browser track depicting CTCF CUT&RUN data at the β-globin locus. Grey: CTCF site with ATAC-seq signal I: Mean mCherry fluorescence of cell lines with *eNanog* at 75 kb or 25 kb with and without deletion of the CTCF motif as indicated J: Representative FACS plots showing an individual clonal cell line with the *Fgf5* E1 enhancer integrated at the 1.5 kb LP. K: Quantification of mean mCherry expression of *Fgf5* E1 integrated at the 1.5 kb LP compared to the no enhancer control. Statistical significance was tested (p < 0.05, one-sided paired t-tests).

**Figure 2:**
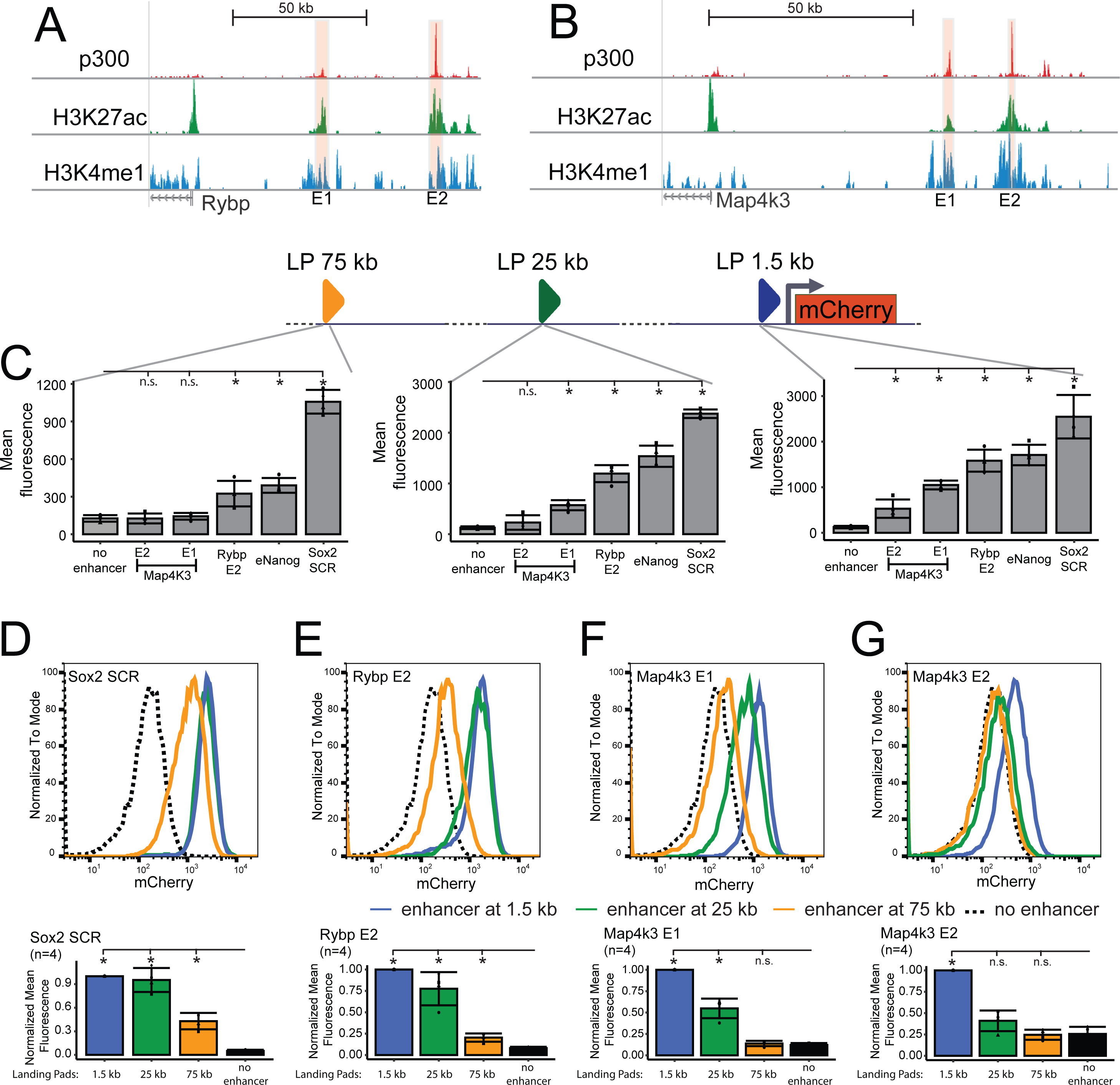
Enhancer strength determines dependency on enhancer-promoter distance. A: UCSC browser track depicting p300, H3K27ac, and H3K4me1 ChIP-seq tracks for the Rybp locus (Buecker et al. 2014) B: UCSC browser track depicting p300, H3K27ac and H3K4me1 ChIP-seq tracks for the Map4k3 locus (Buecker et al. 2014) C: Quantification of mCherry expression of the five indicated enhancers compared to the no enhancer control at three different LPs. Left: integration at 75 kb, middle: integration at 25 kb, Right: integration at 1.5 kb. Statistically significant signals are marked by stars compared to the no enhancer control (p < 0.05, one-sided paired t-tests). n.s.: not significant, p>0.05. Note that for Map3k4 E1 at 75 kb and Map3k4 E2 at 25 kb, additional replicates (see Figures 3 and 4) were performed that decreased p below 0.05. D-G: Quantifying mean mCherry expression of indicated enhancers integrated at the indicated landing pads. Top: individual examples, bottom: normalised mCherry expression. All mean expressions were normalised to the 1.5 kb integration. Statistically significant different signals (p < 0.05, one-sided paired t-tests) are marked by stars.

The fluorescent reporter gene allows for a quick read-out of overall expression levels by FACS. Since mCherry levels are measured in each individual cell, we can obtain information on average expression values and the distribution of expression levels within a population of cells containing the desired enhancer landscape. The final reporter cell lines with the integration of the reporter gene and all three LPs, but without any enhancers, had slightly elevated levels of mCherry fluorescence compared to WT v6.5 cells (Figure 1B clone 1, Figure S1A clone 2).

For integrating enhancers at the reporter locus, we used a variation of the Cre/lox recombination system. Lox sites consist of a spacer sequence that needs to be identical for two lox sites to recombine and two Cre-recognition sites 5’ and 3’ of the spacer. Recombination between one lox site within the genome and one lox site within a donor plasmid leads to the integration of the plasmid into the genome, albeit highly inefficiently since this process is reversible, resulting in the excision of the integrated plasmid. However, the efficiency of plasmid integration can be increased by the use of lox66 and lox71 variants that contain mutations in the left and right Cre-recognition site, respectively (Araki, Araki, and Yamamura 1997; Branda and Dymecki 2004). Their recombination results in one lox site without any mutations and one lox72 site that is no longer recognised by the Cre-recombinase efficiently, as both left and right recognition sequences are mutated; thus, plasmid excision is greatly diminished. We integrated lox71 sites 1.5 kb 5’ of the reporter gene and designed plasmids containing lox66 sites for enhancer integration. Upon recombination between a lox71 site in the genome and a lox66 site in the plasmid, the entire plasmid is integrated into the genome (Fig. 1C). To remove the plasmid backbone after integration, we included two additional compatible FRT recombination sites (Branda and Dymecki 2004) that are orthogonal to the Cre/Lox-system and are flanking the backbone upon plasmid integration (Fig. 1C). Expression of FlpO recombinase leads to recombination between the two FRT sites and excision of the entire plasmid backbone (Fig. 1C), leaving only the enhancer and a single FRT site behind. For the LPs at 25 kb and 75 kb, we chose a similar strategy but used lox variants with different spacer regions that do not recombine with lox71 sites or with each other (here: lox2272-71 and loxm2-71). We designed specific targeting vectors for every landing pad, each with a compatible lox66 site, a particular selection cassette for positive selection fused to ΔTK for negative selection, and an LP-specific set of FRT sites compatible with each other, but not with the FRT sites used for any of the different targeting vectors (see Supplemental Table 6). We validated that all lox sites and integrations occurred as expected for the two independent reporter cell line clones and multiple cell lines with integrated enhancers using Cas9 seq at the β-globin locus in combination with Nanopore sequencing ((Gilpatrick et al. 2020), data not shown).

### The *Nanog* enhancer activates reporter gene expression in a distance-dependent manner

To distinguish the enhancer fragments integrated at the reporter locus from the endogenous enhancers, we amplified all selected enhancers in this study from the *Mus musculus castaneus* strain. As a first proof of concept for enhancer integration, we inserted the proximal enhancer from the *Nanog* locus (Fig. 1D) into a targeting plasmid for the 1.5kb landing pad. After transfection of the enhancer-plasmid along with a plasmid expressing Cre-recombinase and one week of positive selection, roughly 25% of colonies had the desired plasmid integration (data not shown). mCherry expression was slightly decreased upon plasmid integration compared to the reporter cell line without integration (Fig. S1B, right). We then transfected a plasmid expressing FlpO-recombinase and selected for excision of the plasmid backbone with Ganciclovir. In the selected individual clones, mCherry expression was strongly activated, and several independently derived cell lines showed almost indistinguishable mCherry levels (Fig. 1E), demonstrating that our reporter system can measure enhancer activity in a reproducible manner.

Interestingly, some clones initially had a small fraction of mCherry-negative cells that were quickly lost upon passaging (Fig. S1D). To ensure that the mCherry-negative fraction was due to delayed activation of mCherry-expression rather than silencing in mCherry-expressing cells (Cabrera et al. 2022), we sorted mCherry-positive and −negative cells for one of these clones and measured mCherry-fluorescence by FACS every few passages. While the mCherry-positive population stayed positive, the negative population quickly gained mCherry expression (Fig. S1D). This data suggests that initiation of mCherry-expression after the removal of the plasmid backbone can be delayed among individual cells but will be completed within a few passages in the entire population.

Next, we tested the same enhancer at different distances and inserted the *Nanog* enhancer into the targeting plasmids for the LP 25kb and the LP 75kb. Integration efficiencies were similar to integration at 1.5 kb (see Supplemental Table 6 for a summary of integration efficiencies). At both loci, mCherry expression was not affected by plasmid integration (Fig. S1B, middle and left). Upon excision of the backbone, we selected several independent clones with the *Nanog* enhancer integrated at 25 kb or 75 kb. The different clones with integration at 25 or 75 kb behaved very similarly (Fig. 1E, middle and left). We did not detect a fraction of mCherry-negative cells with the enhancer at either 25 or 75 kb, and expression levels were stable over several passages (data not shown). Next, we compared mCherry expression across different *Nanog* integration distances (Fig. 1F, 1G). We calculated the mean fluorescence and subtracted the mean background autofluorescence of the unmodified WT v6.5 control (Fig. 1B) to compare the activation of mCherry across replicates. In a previous study, the levels of reporter gene fluorescence were linearly correlated with the number of mRNAs as determined by single-molecule RNA FISH (Zuin et al. 2022). Therefore, we used the background-subtracted mean fluorescence as a direct read-out of the transcriptional activity of the reporter gene. We normalised the background-subtracted mean fluorescence at each LP to the background-subtracted mean fluorescence at 1.5 kb to compare reduced activity at larger genomic distances to the initial intrinsic strength of the enhancer at 1.5 kb (Figure 1G and S1C). The expression of mCherry is slightly decreased upon integrating the *Nanog* enhancer at 25 kb compared to 1.5 kb. However, when the enhancer is moved to the 75 kb landing pad, mCherry expression is strongly decreased across the population but is still higher than the no enhancer control. In summary, our reporter system allows for the efficient integration of enhancer sequences at three different distances from a reporter gene and confirms the recently reported importance of genomic enhancer-promoter distance (Zuin et al. 2022; Rinzema et al. 2022).

CTCF sites can interfere with enhancer-promoter communication (Zuin et al. 2022; de Wit et al. 2015; Pachano, Haro, and Rada-Iglesias 2022) and could be responsible for the loss of activity of the *Nanog* enhancer at 75 kb. We analysed multiple publicly available CTCF cut&run datasets (Patty and Hainer 2021; Olbrich et al. 2021) and identified one CTCF peaks between the *Hbb-bt* and the *Olfr67* gene, i.e., between the 25 kb and the 75 kb LPs (Figure 1H). This peak was accessible in ATAC-seq experiments performed on the parental reporter cell line (Figure S1E), suggesting that CTCF might indeed bind it. Therefore, we asked whether the CTCF binding site affects the communication of enhancers integrated at 75 kb with the mCherry promoter. We deleted the CTCF site with guides flanking the CTCF motifs in cell lines where the *Nanog* enhancer was incorporated at the 75 kb landing pad. Next, we compared mCherry expression between the cell lines with and without the CTCF motif: mCherry expression was slightly increased in those cells where the CTCF motif was deleted (Figure 1I). However, this increase is relatively minor compared to the Nanog enhancer at 25 kb. We conclude that CTCF binding between the 25 and 75 kb LP slightly reduces mCherry expression and is partially responsible for the drop in mCherry expression when integrating an enhancer at 75 rather than at 25 kb. However, the increased distance is primarily responsible for the reduction in mCherry expression.

Next, we wanted to test whether increased mCherry expression is due to the integration of active enhancers or whether the reporter gene can be activated by integrating any DNA sequence. Therefore, we integrated the *Fgf5* E1 enhancer in the 1.5 kb LP. *Fgf5* E1 is only activated during the differentiation of mouse ESCs into so-called EpiLCs, is not actively repressed in mouse ES cells and has very low intrinsic strength in luciferase assays even when active (Thomas et al. 2021). As expected, integration of the *Fgf5* E1 enhancer at 1.5 kb did not increase mCherry expression beyond the no enhancer control (Figure 1J, 1K and S1F).

The β-globin locus is inactive in mouse ESCs; however, individual promoters might be activated by integrating the different enhancers. We performed ATAC-seq in selected cell lines with different integrated enhancers. We analysed the open chromatin landscape at the β-globin locus and the *Nanog* locus with and without the enhancers and in the parental cell lines (Figure S2). No differences were observed between the cell lines, suggesting that the manipulations did not dramatically change the cell identity and that the minimal TK promoter is the only promoter activated at the synthetic locus.

In summary, we have generated a highly versatile synthetic locus that allows for the reproducible interrogation of enhancer activity at defined distances to a promoter controlling the expression of a reporter gene. The locus itself is inert, expression of the reporter gene is not silenced over time and reporter gene expression is only activated by active enhancers

### Enhancer strength determines dependency on enhancer-promoter distance

Recently, both the SCR from the *Sox2* locus in ESCs and the human β-globin micro-LCR in erythroleukemia K562 cells have been demonstrated to depend on the genomic distance to the promoter of a reporter gene in a model locus (Rinzema et al. 2022; Zuin et al. 2022). In the case of the SCR, measuring expression levels systematically at many different genomic distances revealed that target gene expression decays non-linearly with increased distance (Zuin et al. 2022). However, both studies relied on a highly active enhancer, so whether all enhancers respond to genomic distance in the same way is currently unknown. The high efficiency of enhancer integration into our reporter locus enabled us to systematically compare how different enhancers are affected by genomic distance. We chose enhancer elements that were part of multi-enhancer clusters and showed different strengths in a previously published STARR-seq data set (Peng et al. 2020). We included the proximal *Nanog* enhancer (see Figure 1 (Agrawal et al. 2021)), the E2 enhancer element located about 90 kb upstream of the *Rybp* locus (Fig. 2A) and two enhancers (E1 and E2) from the *Map4k3* locus (60 and 75 kb upstream of the transcription start site (TSS), Fig. 2B). We first tested the ability of these four enhancers to activate expression in a plasmid-based luciferase assay. All enhancers significantly activated luciferase activity (Fig. S3A), albeit to different degrees: The *Nanog* and *Rybp E2* enhancers were the most potent activators, followed by *Map4k3 E2* and finally *Map4k3 E1*. In addition, we included the well-studied Sox2 control region (SCR), a 6 kb multi-enhancer cluster roughly 100 kb upstream of the *Sox2* gene (Zuin et al. 2022; Taylor et al. 2022).

Next, we integrated every enhancer into each landing pad of the reporter locus individually and determined the resulting mCherry levels by FACS (clone 1: Figure 2C, clone 2: Figure S3B). Integration of any selected enhancer at 1.5 kb increased mCherry expression levels (Fig. 2C, right, S3C right), showing that all tested elements act as enhancers and can activate transcription. The *Sox2* SCR was the strongest activator, followed by the *Nanog* and *Rybp E2* enhancers, *Map4k3 E1* and *Map4k3 E2*. The expression levels of mCherry were reproducible between different clones and replicates, but variability is increased for the lower expressing cell lines with *Map4k3 E1* or *Map4k3 E2* integration. Of note, despite higher activity in luciferase assays, mCherry levels upon integration of *Map4k3 E2* were lower compared to integration of *Map4k3 E1* (Fig. 2C). Next, we analysed the expression of mCherry when the same enhancers were integrated at either 25kb (Figure 2C middle, S3B middle) or 75 kb (Figure 2C left, S3B left). At all distances, the order of enhancer strength was preserved: *Sox2 SCR* activated the highest levels of mCherry expression at all distances, followed by *eNanog*, *Rybp E2, Map4k3 E1* and finally *Map4k3 E2*.

While the order in enhancer strength was preserved, the activation of mCherry expression dropped with increasing distances (clone 1: Figure 2C, clone 2: S3B). The strong enhancers *Sox2 SCR* (Figure 2D, S3C), *eNanog* (Figure 1G, S1C) and *Rybp E2* (Figure 2E, S3D) showed strong activation at 1.5 kb and 25kb, with no (*SCR*) or only slight loss (*eNanog* and *Rybp E2*) of activity at the intermediate distance 25 kb. When these strong enhancers are integrated at 75kb, mCherry expression is substantially decreased but still detectable and much stronger than the no enhancer control. For the two weaker enhancers, *Map4k3 E1* (Figure 2F, S3E) and *Map4k3 E2* (Figure 2G, S4F), the activation from 25 kb is strongly reduced and almost undetectable at 75 kb. Of note, in the first set of experiments shown in Figure 2, neither *Map4k3 E1* at 75kb (Figure 2C, 2F) nor *Map4k3 E2* at 25kb (Figure 2C, 2G) in clone 1 reached statistical significance. However, with increasing replicates (see Figures 3B-E and 4A-D), both enhancers reached statistical significance at these loci. Nevertheless, the measured activity is very low, and the relative loss of activity compared to 1.5 kb is much more pronounced than for the stronger enhancers at the same distance (Figure 2C, S3B). Together, these experiments show that increasing distance to the minimal promoter decreases the ability of a selected enhancer to activate that promoter. Still, stronger enhancers are much less affected by a similar increase in genomic distance and can activate this promoter from larger distances than weaker enhancers.

**Figure 3:**
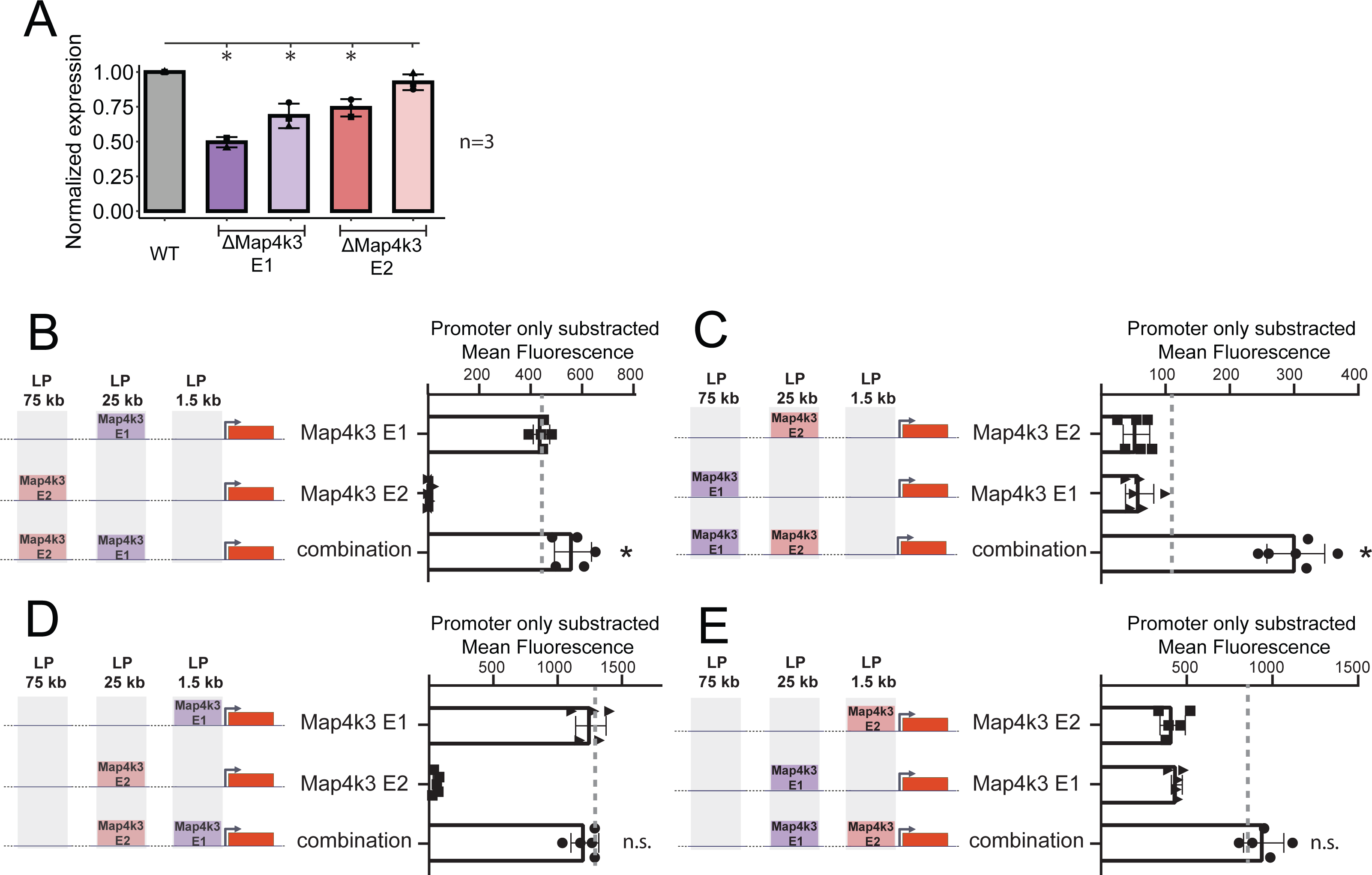
Weak enhancers activate transcription synergistically from a distance. A: Quantification of Map4k3 expression after deletion of the two different individual enhancers compared to the WT expression. B-E: Comparison of combinations of enhancers to the individual integrations and the expected additive behaviour. Left: schematic of individual and combinations of integrations at the different LPs. The mean mCherry expression was calculated, and then the mean expression of the no-enhancer control was subtracted from all individual and combination mean mCherry expressions. The Grey dashed line indicates the expected values under an additive model. Statistically significant signals compared to the expected additive modell (p < 0.05, one-sided paired t-tests) are marked by stars or n.s. (not significant). B: *Map4K3 E2* at 75 kb combined with *Map4K3 E1* at 25 kb C: *Map4K3 E1* at 75 kb combined with *Map4K3 E2* at 25 kb D: *Map4K3 E2* at 25 kb combined with *Map4K3 E1* at 1.5 kb E: *Map4K3 E1* at 25 kb combined with *Map4K3 E2* at 1.5 kb

### The cooperation of weak enhancers can activate transcription from a distance

Both enhancers from the *Map4k3* cluster were the weakest elements tested above; specifically, could not elicit considerable mCherry expression from 25 kb. At the endogenous locus, this element is located about 75 kb from the potential target gene and, therefore has to overcome a substantially longer distance to the promoter. Consequently, we tested whether *Map4k3 E2* contributed to the expression of the putative target gene *Map4k3*. We deleted both *Map4k3* enhancers individually at the endogenous locus using CRISPR-Cas9. We selected two clones each and analysed the expression of the target gene *Map4k3* using qPCR. Deletion of *Map4k3 E1* clearly reduced *Map4k3* expression in both analysed clones (Figure 3 A). Deletion of the *Map4k3 E2* enhancer also reduced the expression of the target gene (Figure 3A), albeit not as strongly as the Map4k3 E1 enhancer and only in one of the two tested clones.

As the weak enhancer element *Map4k3 E2* affects expression levels at the endogenous locus from a distance at which it is not active in our synthetic locus, we hypothesised that the presence of additional enhancers influences how a given enhancer is affected by genomic distance. Therefore, we integrated both *Map4k3 E1* and *E2* elements at different distances into our synthetic locus. As the simultaneous integration of two enhancers was very inefficient, we performed the dual integrations step-wise: we integrated one enhancer first, selected individual clones, and then integrated the second enhancer. Both backbones were removed after the second enhancer was integrated. As the two parental cell line clones behaved very similarly upon integrating individual enhancers (Fig. 1 and 2), we used only clone 1 for all dual enhancer experiments described below. In the previous experiments described in Figure 2, we compared the integrated enhancers to the no-enhancer control. In the following experiments, we compare dual integrations to individual integrations, and we calculate values of expected activity for enhancer combinations based on an additive model of their individual activities (gray dotted lines in Fig. 3B-E and 4A-D). To avoid accounting for basic expression levels from the promoter twice when adding up the auto-fluorescence corrected expression values of two individual enhancer integrations, we subtracted the background mCherry levels of the no enhancer control from the measured mean mCherry expression.

First, we tested whether *Map4k3 E2* at 75 kb can increase expression levels in the presence of the *Map4k3 E1* enhancer at 25kb. Interestingly, the combination of the two elements lead to an increased expression of mCherry compared to the individual integrations (Figure 3B), even though *Map4k3 E2* cannot activate the reporter gene from 75 kb by itself. Next, we swapped Map4k3 E1 and Map4k3 E2 positions: now *Map4k3 E1* was integrated at 75 kb, and *Map4k3 E2* was integrated at 25 kb. Both enhancers show very low individual activation of mCherry fluorescence at these positions; however, in combination, they strongly increase mCherry expression (Figure 3C) and activate transcription super-additively. We conclude that weak enhancers can work synergistically, allowing combinations of two enhancers to activate transcription from distances where the individual enhancers are not active.

In the experiments above, we integrated the enhancers at the two distal locations. Next, we moved the two elements closer to the promoter and integrated *Map4k3 E1* and *E2* at the 1.5 and 25 kb positions. First, we tested the stronger enhancer (*Map4k3 E1*) at 1.5 kb and found that integration of the weaker enhancer (*Map4k3 E2*) at 25 kb did not further increase the expression of mCherry (Figure 3D). When we switched those positions and integrated the weaker element *Map4k3 E2* at 1.5 kb and *Map4k3 E1* at 25 kb (Figure 3E), the combined integration increased the expression of mCherry. However, it increased only slightly beyond the expected additive behaviour. In both cases, the minimal promoter itself may be a limiting factor in the combined action of the two elements and saturation of the promoter might not allow for an additional increase in activation (see discussion)

### Cooperation between strong and weak enhancers depends on their relative position

*Map4k3 E2* and *Map4k3 E1* belong to the same enhancer cluster and can cooperate. We next asked whether the ability to cooperate with a second enhancer is limited to elements from the same locus or whether the weak *Map4k3 E2* enhancer can also work together with other enhancers. We first combined *Map4k3 E2* with the *Nanog* enhancer in our synthetic locus. When the *Nanog* enhancer was integrated at 25kb, adding *Map4k3 E2* at 75 kb did not increase the expression of mCherry further (Figure 4A). However, when we changed the order and integrated the *Nanog* enhancer at 75 kb and *Map4k3 E2* at 25kb, we observed a strong super-additive induction of mCherry expression compared to the individual integrations (Figure 4B). This synergistic effect was not limited to the *Nanog* enhancer: when we instead integrated *Rybp E2* at 75 kb together with *Map4k3 E2* at 25kb, the expression of mCherry also strongly exceeded the expected additive values (Figure 4C).

**Figure 4:**
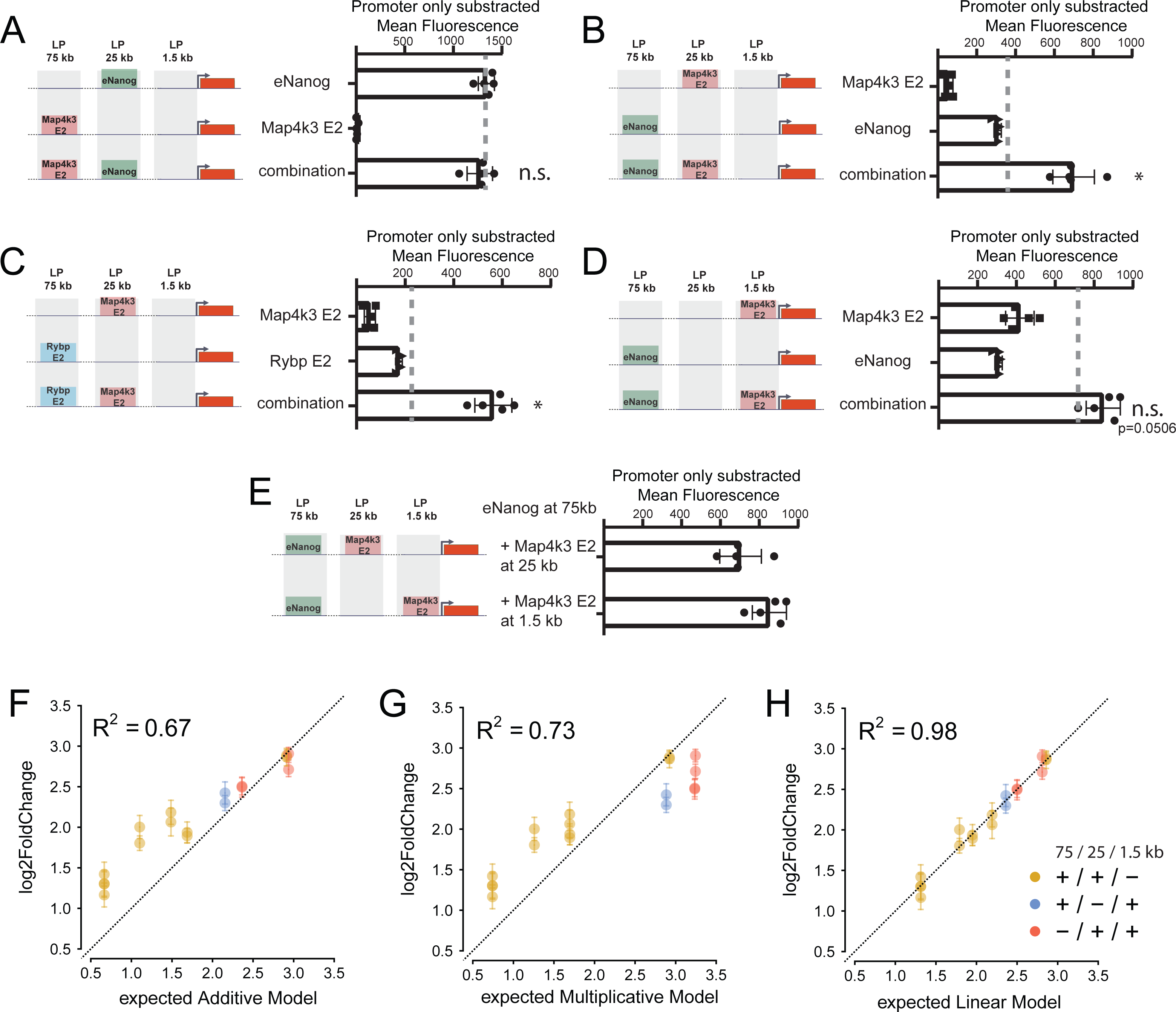
*Map4k3 E2* can cooperate with strong enhancers to activate transcription from a distance. A-D: Comparison of combinations of enhancers (third row) to the individual integrations (top two rows) and the expected additive behaviour. Left: schematic of individual and combinations of integrations at the different LPs. The mean mCherry expression was calculated, and then the mean expression of the no-enhancer control was subtracted from all individual and combination mean mCherry expressions. The Grey dashed line indicates the expected values under an additive model. Statistically significant signals compared to the expected additive modell (p < 0.05, one-sided paired t-tests) are marked by stars or n.s. (not significant). A: *Map4K3 E2* at 75 kb combined with *eNanog* at 25 kb B: *eNanog* at 75 kb combined with *Map4K3 E2* at 25 kb D*: Rybp E2* at 75 kb combined with *Map4K3 E2* at 25 kb E: eNanog at 75 kb combined with *Map4K3 E2* at 1.5 kb E: Comparison of background subtracted mean mCherry expression of *eNanog* at 75 kb and *Map4k3 E2* at 25 kb (top) or 1.5 kb (bottom) F: Additive model of enhancer activity at the different loci G: Multiplicative model of enhancer activity at the different loci H: Linear Regression Model to predict enhancer activity

Thus far, we have shown that *Map4k3 E2* can support the distal enhancers when integrated at the intermediate LP at 25kb. Next, we asked whether this synergistic effect is strengthened when integrated closer to the promoter. Therefore, we moved *Map4k3 E2* to the 1.5 kb position in a cell line where the *Nanog* enhancer was integrated at 75 kb. Combined enhancers in these cell lines lead to increased mCherry expression compared to single enhancers (Figure 4D). However, the overall expression levels were only slightly above the added single enhancer levels. We compared the mCherry expression levels between cell lines where the *Nanog* enhancer is integrated at 75 kb and *Map4K3 E2* is either located at 25 kb or 1.5 kb (Figure 4E). While the expression of mCherry is higher in cell lines with *Map4k3 E2* at 1.5kb, this is only a minor increase compared to *Map4k3 E2* at 25kb. In summary, we conclude that a weak enhancer such as *Map4k3 E2* can synergise with strong enhancers when integrated between the promoter and the strong enhancer, thereby partially alleviating the distance-dependent drop in activation of the strong enhancer. This behaviour does not strongly depend on the exact position where the weak *Map4k3 E2* enhancer is integrated. We conclude that for expression levels of the target gene, the position of the weaker enhancer in relation to the stronger enhancer is flexible.

Two recent publications have analysed enhancer synergies using an episomal plasmid approach but reached different conclusions (Loubiere et al. 2023; Martinez-Ara, Comoglio, and Steensel 2023): In mouse ES cells, the enhancers worked predominantly additive (Martinez-Ara, Comoglio, and Steensel 2023), whereas the work conducted in Drosophila S2 cells showed that most developmental enhancers are synergistic and follow a simple multiplicative model, with the caveat that strong enhancers might saturate limited promoter capacity (Loubiere et al. 2023). We examined whether either model could explain our observations: The simple additive model assumes that summing the individual activities of two enhancers at their respective distance can predict their combined activity (see Methods). For combinations where one of the enhancers is integrated proximal to the promoter, we found that this model describes our data very well (Figure 4F). However, the distal combinations, except for *eNanog* at 25kb and *Map4k3 E2* at 75 kb, do not follow this trend and show super-additive behaviour (Figure 4F). Next, we tested a multiplicative model, but this model described none of our combinations well (Figure 4G). While the activity of distal enhancer combinations was higher than expected for multiplicative behaviour, the combinations with a proximal enhancer had lower than expected activity (Figure 4G). Because neither the additive nor the multiplicative model takes the position of the enhancers relative to the promoter into consideration, we instead fitted a linear regression model that factors in the distance at which the two cooperating enhancers are integrated to predict their combined activity from their individual contributions. The linear model improved the prediction accuracy strongly (R²= 0.98, Fig. 4H), suggesting that including information about the distance at which the cooperating enhancers are integrated is required and sufficient to predict the activity of enhancer combinations accurately. One key asset of linear models is to deliver a set of interpretable and informative coefficients. The individual activities at the different positions are most predictive of the combined activity. In addition, the model penalises combinations where an enhancer is localised at 1.5 kb. Furthermore, it predicts that combinations of enhancers at 25 and 75 kb have a strong synergistic effect (see supplementary table 7 for the full report). In conclusion, how two enhancers cooperate depends strongly on their genomic distance to the promoter: While enhancers integrated at 25 and 75 kb synergise strongly, a simple additive model is sufficient to describe the data, when one enhancer is integrated close to the promoter. Furthermore, similar to previous studies (Loubiere et al. 2023), saturation of the promoter might be limiting more substantial synergistic effects.

## Discussion

Here, we developed a platform for studying distance-dependent enhancer-enhancer cooperativity in the mammalian genome. The platform includes a reporter cell line for integrating enhancer sequences at three different distances upstream of a reporter gene to ask how individual enhancers and their combinations activate transcription from different genomic distances in a standardised genomic environment. Our system has several advantages over existing approaches. It is very flexible, as we can integrate any sequence and combinations at currently three different positions of the reporter locus. As the locus contains no active regulatory elements besides the ones we are introducing, it provides a controlled environment for building and analysing complex regulatory landscapes de novo. Furthermore, enhancers are tested in their native chromatinized environment, while the efficiency of our system allows us to generate multiple cell lines at once and test more combinations of regulatory elements than would be possible by modifying endogenous loci.

To demonstrate the power of the platform and reporter cell line, we integrated five different enhancers at 1.5, 25 and 75 kb upstream of a mCherry reporter gene and measured the resulting fluorescence. mCherry levels were very similar between independently generated clonal cell lines. Interestingly, plasmid integration at 1.5 kb but not at 25 or 75 kb reduced mCherry expression before FlpO treatment. Even after FlpO-expression, a small fraction of mCherry-negative cells remained in some clones with e*Nanog* integration at 1.5 but were lost after a few passages. We did not observe a similar behaviour at 25 or 75 kb. We speculate that upon plasmid integration, read-through transcription of the selection cassette reaches the minimal TK-promoter of the mCherry reporter gene and leads to deposition of H3K36me/me2/me3, which in turn recruits the DNA methylation machinery (Weinberg et al. 2019). Silencing by DNA methylation could explain reduced mCherry expression upon plasmid integration. After excision of the selection cassette, the enhancer would then activate the reporter gene and DNA methylation is removed. The persistence of DNA methylation for some time could explain the small fraction of mCherry-negative cells that is lost upon passaging cells.

We have built the synthetic locus around the weak minimal TK promoter that shows very low basal activity. This ensures that the integrated enhancers indeed cause all measured activity at the locus. However, this promoter might also be rate limiting in our analysis: Most strong enhancers and combinations at close distances showed similar mean expression levels, suggesting saturation in expression. However, the 6 kb multi-enhancer from the *Sox2* SCR showed much stronger expression than the other enhancers, even from the 25 kb distance, suggesting that stronger expression is possible. Here, combinations of very strong enhancers and the incorporation of different promoters in the future will test the limitations of the system.

All enhancers tested in this study depend on the genomic distance to their target promoter, albeit to different degrees according to their intrinsic enhancer strength. Our results agree with and expand on previous findings regarding the distance dependency of two highly active regulatory regions in mouse ESCs and erythroleukemia K562 cells (Zuin et al. 2022; Rinzema et al. 2022). While all enhancers lost activation potential with increasing distance, introducing a second, weak enhancer can partially rescue the drop in activation. We focused on one such weak enhancer, the *E2* enhancer from the *Map4k3* locus. In isolation, this element cannot activate transcription even from intermediate distances. However, when combined with other enhancers and placed between the strong enhancer and the promoter, this element strongly increased the activation at the promoter.

While increasing the distance between enhancer and promoter decreases the activation at the promoter, many enhancers are located at distances where the individual enhancers should not be able to activate a promoter anymore. Our work here suggests that integrating additional, weak enhancers can partially overcome loss of enhancer activity due to increased genomic distance. Interestingly, the individual intrinsic strength of the additional enhancer and its exact position might be secondary as long as it is located between the strong enhancer and the promoter. Our findings might explain recent findings from the ɑ-globin (Blayney et al. 2022) and the *Sox2* loci (Brosh et al. 2023), where elements with weak intrinsic enhancer activity can facilitate the communication of strong enhancers with the promoter, even though we cannot rule out that some of these elements apply distinct mechanisms of enhancer cooperation.

Why can stronger enhancers bridge more considerable genomic distance than weaker ones and how do weak enhancers synergize with distal strong enhancers? Increased genomic distance reduces contact frequency (Dixon et al. 2012; Rao et al. 2014). However, enhancer-promoter contacts do not linearly correlate with transcriptional activation of the target gene (Zuin et al. 2022), and it remains unclear whether and how close an enhancer needs to come to its promoter to activate transcription (H. Chen et al. 2018; Benabdallah et al. 2019; Alexander et al. 2019). Regulatory elements recruit tens of TFs, co-factors and RNA Pol II molecules (Li et al. 2019; 2020). These so-called transcription hubs of regulatory factors can be several 100 nm in size and might be able to bridge enhancers to their target promoters (Heist, Fukaya, and Levine 2019). In that case, the actual 3D distance between an enhancer and a promoter would not matter as much as long as the transcription hub still “touches” the promoter. We speculate that stronger enhancers have more prominent transcription hubs due to the recruitment of higher levels of regulatory factors and might thus be capable of activating their target gene from a larger 3D distance. This would increase resilience to small increases in 3D distance and thus decrease dependence on genomic enhancer-promoter distance. Such a model could explain why the *Sox2 SCR* activates almost identical expression levels from 1.5 and 25 kb, whereas the weaker enhancers are much less active from 25 kb compared to 1.5 kb. Similarly, weak enhancers at intermediate distances could bridge the transcription hubs of strong distal enhancers to the promoter. They thereby increase the activation radius of the strong enhancer elements.

In previous work, we demonstrated that elements without intrinsic strength measured in reporter activity can contribute to gene expression. Here, we expand on this study and show that weak enhancers can strongly contribute to the expression of a target gene, potentially by facilitating the communication of the enhancer with the promoter. It is possible that the mere positioning between the strong enhancer and the promoter could be enough to bridge the distance between the distal enhancer and the promoter. A recent MPRA approach in different human cell lines classified cis-regulatory elements into different categories based on their intrinsic activity, including classical enhancers and so-called chromatin-dependent enhancers. The latter category is characterised by a strong epigenomic enhancer signature but low intrinsic enhancer strength. Enhancers from both categories can often be found within the same enhancer cluster. The combination of weak and strong enhancers in the same enhancer cluster could fulfil multiple functions. On the one hand, multiple weak enhancers might confer a buffering effect against the potential loss of individual elements. In addition, our results suggest that combining enhancers with weak and strong intrinsic activity can expand the genomic distance at which an enhancer cluster can communicate with its target gene. Moreover, we speculate that enhancer cooperativity could improve efficiency in target promoter selection: promoters associated with an additional weak enhancer may exhibit an increased responsiveness to a distal strong enhancer compared to other promoters without such an element.

## Acknowledgements

We would like to thank all members of the Buecker lab for discussions and feedback throughout the project, Martin Leeb, and the members of his lab for continuous support and critical feedback in shared group meetings. In addition, we would like to thank Patricia Rothe and Mustafa Alaboo for technical help. The BioOptics-FACS facility at the Max Perutz labs was instrumental in the success of this project. Cas9-seq was performed by the Next Generation Sequencing facility at the Vienna BioCenter Core Facilities (VBCF), a member of the Vienna BioCenter (VBC), Austria. Some figures were created with BioRender. The work was supported by the Austrian Science Fund FWF (P34123 to CB and W1261 DK SMICH) and Uni:Docs fellowship from the University of Vienna to H.T.

## Figure Legends

## Methods

### Generation of reporter cell line

All generated cell lines and plasmids are summarized in Supplemental Table 1, all gRNA sequences can be found in Supplemental Table 2, all primers for cloning can be found in Supplemental Table 3 and the sequences for the ssDNA oligos are listed in Supplemental Table 5. Finally, qPCR primers are listed in Supplemental Table 4.

For KI of the reporter gene into the β-globin locus, we first generated a gRNA-expressing plasmid and a targeting vector. Therefore, a single gRNA was designed to target the locus roughly 1.5 kb downstream of the *Hbb-y* gene. As described before, the gRNA was cloned into Addgene plasmid #42230 (Thomas et al. 2021). Due to a SNP in the underlying sequence, this gRNA recognises the C57BL/6 but not the 129/sv allele of v6.5 mouse ESCs, allowing us to introduce the reporter gene specifically into the β-globin-locus on the C57BL/6 allele.

We used Gibson assembly to generate the targeting vector to insert the reporter gene into the β- globin-locus unless noted otherwise. Homology arms were designed to disrupt the gRNA recognition sequence upon successful KI.

We first amplified a 77 bp long minimal TK promoter followed by the mCherry coding sequence by PCR and inserted the fragment into an NcoI-HF®-digested (NEB) pGemT-vector (Promega). We amplified the left homology arm (637 bp) targeting the β-globin locus roughly 1.5 kb downstream of the *Hbb-y* gene from R1 ESC DNA by PCR. We inserted it upstream of the TK-promoter into the SphI-HF®-digested (NEB) TK-mCherry plasmid, leaving the SphI-site intact. We used this SphI site to introduce a loxP-flanked puromycin-deltaTK cassette and a 1.5 kb spacer from inert human DNA between the left homology arm and the TK promoter. The loxP-flanked puromycin-deltaTK cassette was PCR-amplified from a previously generated targeting vector (Thomas et al. 2021). Using a forward primer that mapped to the loxP site upstream of the puromycin-deltaTK cassette but contained mutations, we turned the upstream loxP site into a lox71 site. The 1.5 kb spacer was amplified from HEK-293 DNA, kindly provided by the lab of Dea Slade. We then inserted a 232 bp long BGH polyA sequence downstream of the mCherry coding sequence into the NotI-HF®-digested (NEB) plasmid, leaving the NotI-site intact and introducing an additional BamHI-site. As we did not manage to PCR-amplify the right homology arm, we ordered the synthesised fragment (832 bp) integrated into a plasmid and flanked by a BamHI- (upstream) and a NotI-site (downstream) from BioCat GmbH. We obtained the right homology arm fragment by BamHI-HF®- and NotI-HF®-digestion (both NEB) and ligated it into the likewise BamHI-HF®- and NotI-HF®-digested HAL-pdTK-spacer-TK-mCherry-pA plasmid, downstream of the polyA site.

For KI of the reporter gene, v6.5 mouse ESCs^38^ were cultured and transfected by lipofection with 400 ng of circular targeting vector, 400 ng of gRNA-containing plasmid and 4 μL of Lipofectamine 2000 Transfection Reagent (Invitrogen^TM^), as described before (Thomas et al. 2021). In brief, cells were transferred to a 10 cm dish the day after transfection and selection with Puromycin (2 μg/mL, InvivoGen) was started within 48 h of transfection. After a week of selection, single colonies were picked. Integration was validated by PCR with primers “KI validation 1 forward+reverse” (mapping upstream of the left homology arm in the genome and downstream of the left homology arm in the insert) and “KI validation 2 forward+reverse” (mapping upstream of the right homology arm in the insert and downstream of the right homology arm in the genome). All analysed clones were heterozygous, with the reporter gene being inserted on the C57BL/6 allele as judged by SNPs in Sanger-sequenced PCR products. Clones were expanded and subsequently transfected with a plasmid expressing Cre-recombinase to remove the puromycin-deltaTK cassette and leave a single lox71 site (in reverse orientation) behind. After one week of selection with Ganciclovir (500 ng/mL, InvivoGen), single clones were picked, and removal of the puromycin-deltaTK cassette was confirmed with primers “KI validation after Cre forward+reverse” (mapping up-and downstream of the excised cassette). Overlapping PCR products (primers “reporter validation 1-4 forward+reverse”) spanning the entire insert, the homology arms and the genome are up- and downstream of the insertion were submitted for Sanger sequencing to confirm the intactness of the β-globin-locus, the homology arms and the inserted sequence.

To introduce lox2272/71 and loxm2/71 sites roughly 25 and 75 kb upstream of the mCherry reporter gene, we designed gRNAs and cloned them into Addgene plasmid #42230. We also ordered single-stranded (ss) DNA oligos from Microsynth AG, containing the respective lox site (34 bp; in reverse orientation) and left and right homology arm. The homology arm at the 5’ end of the respective oligo was 66 bp long, and the homology arm at the 3’ end was 50 bp (150 bp total, including the lox site). Homology arms were designed to disrupt the gRNA recognition sequence upon successful KI.

We used two independent clones with reporter gene integration to sequentially knock-in the two lox sites. Therefore, we transfected 500 ng of gRNA-expressing plasmid, 1000 ng of ssDNA-oligo and 100 ng of plasmid expressing a fluorescent marker with 10 μl of Lipofectamine 2000 Transfection Reagent (Invitrogen^TM^). Transfected cells were processed as described before (Thomas et al. 2021), i.e., single fluorescent cells were sorted into 96-well plates. Successful insertion of the lox sites into the genome was initially confirmed by PCR with forward primers overlapping the integrated lox sites (and thus only binding to the modified alleles) and reverse primers downstream of the integration site (“lox KI 25/75 kb initial forward+reverse”). Selected clones were tested further with a more upstream forward primer binding to both alleles (“lox KI 25/75 kb validation forward”), and resulting PCR products were subcloned into a pGEM-T vector (Promega, pGEM-T Vector Systems) for subsequent Sanger sequencing of both alleles. The recognition sequence of the gRNA used for the lox site at 25 kb contains SNPs on the 129/sv allele, allowing us to integrate the lox2272/71 site on the C57BL/6 allele specifically. For integration at 75 kb, we did not design a gRNA overlapping a SNP. Instead, we chose heterozygous clones with the integration of the loxm2/71 site on the C57BL/6 allele, as judged by SNPs surrounding the integration site.

### Design of plasmids for enhancer integration

We generated three plasmids to integrate enhancers at 1.5, 25 and 75 kb distance upstream of the reporter gene by targeting the lox71, loxm2/71 and lox2272/71 sites. We first introduced a single FRT site into a pGemT-vector (Promega). Therefore, forward and reverse DNA oligonucleotides - containing the FRT sequence as well as the overhangs required for cloning - were ordered from Microsynth AG, annealed and cloned into a NcoI-HF®- and SphI-HF-® digested pGemT plasmid, as described before (Thomas et al. 2021). We generated three plasmids containing FRT, FRT3 and FRT5 sites, respectively. The original restriction sites were disrupted during this process, but an additional NcoI-site was introduced downstream of the FRT sites as part of the inserted oligo. We then ordered additional DNA oligos and introduced lox66 sites downstream of the FRT sites of the resulting NcoI-HF®- and NotI-HF®-digested plasmids (both NEB). For the plasmid containing an FRT site, we introduced a lox66 site; for the plasmid containing the FRT3 site, we introduced a loxm2/66 site; and for the plasmid with the FRT5 site, a lox2272/66 site.

We digested the resulting plasmids by NotI-HF® (NEB) and inserted cassettes expressing fusions of antibiotic resistance genes with deltaTK downstream of the lox66 sites by Gibson assembly. We inserted blasticidin-deltaTK into the FRT-lox66-plasmid, puromycin-deltaTK into the FRT3- loxm2/66-plasmid and hygromycin-deltaTK into the FRT5-lox2272/66-plasmid. The forward primer used for amplification of the cassettes included an additional FRT site in the case of blasticidin-deltaTK, an additional FRT3 site in the case of hygromycin-deltaTK and an additional FRT5 site for puromycin-deltaTK. The NotI site between the lox and the newly inserted FRT site was restored and used to integrate enhancer sequences.

All in all, we generated the following three plasmids:

● pGemT-FRT-lox66-FRT-blasticidin-deltaTK
● pGemT-FRT3-loxm2/66-FRT3-puromycin-deltaTK and
● pGemT-FRT5-lox2272/66-FRT5-hygromycin-deltaTK.

These plasmids were digested by NotI-HF®, and enhancer sequences amplified from *castaneus* mouse strain DNA (kindly provided by the lab of Kikuë Tachibana) were inserted by Gibson assembly. Primers for amplifying enhancer sequences were designed to encompass the central p300 peak (ChIP-seq from Buecker *et al*. (Buecker et al. 2014)). The resulting plasmids were transfected together with a Cre-recombinase expressing plasmid into the reporter cell line for insertion of enhancers at the lox71 sites of the reporter locus.

### Integration of enhancer sequences

To integrate enhancer sequences, 200,000 cells of the reporter cell line were plated and transfected by lipofection on the following day. Therefore, 250 ng of enhancer-containing plasmid, 1750 ng of Cre-recombinase expressing plasmid and 10 μL of Lipofectamine 2000 Transfection Reagent (Invitrogen^TM^) were used. Cells were selected with Blasticidin (10 μg/mL, Invivogen), Hygromycin B (400 μg/mL, Sigma-Aldrich) or Puromycin (2 μg/mL, InvivoGen), depending on the selection cassette present in the enhancer plasmid. Single colonies were picked after a week of selection. Plasmid integration was validated by PCR with forward primers mapping to the integrated plasmid backbone (“integration forward”) and reverse primers mapping downstream of the respective lox site (“integration 1.5/75 reverse; for 25 kb: “lox KI 25 kb initial reverse” that was used before for validating KI of the lox site). Colonies with integration of enhancer plasmid were expanded and transfected with plasmid expressing FlpO-recombinase (a more active, codon-optimized version of Flp kindly provided by the lab of Stefan Ameres) to remove the plasmid backbone including the selection cassette (200,000 cells, 2 μg of FlpO-recombinase expressing plasmid, 10 μL of Lipofectamine 2000 Transfection Reagent (Invitrogen^TM^)). Cells were passaged, seeded at low density the day after transfection and selected with Ganciclovir (5 μg/mL, Invivogen) for one week. Single colonies were picked. PCR confirmed the removal of the selection cassette with an enhancer-specific forward primer combined with a primer downstream of the respective lox71 site (“integration 1.5/75 reverse”; for 25 kb: “lox KI 25 kb initial reverse”). In addition, the PCR with primers “integration forward” and “integration 1.5/75 reverse” or “lox KI 25 kb initial reverse” was repeated. As the forward primer maps to the plasmid backbone, the absence of a band in that PCR confirms the complete removal of the plasmid backbone in the entire cell population.

Allele-specific primers up and downstream of the lox sites were designed. In the case of the lox71 site at 1.5 kb, a forward primer upstream of the lox71 site (“1.5 forward”, recognising both alleles) was combined with a reverse primer mapping to the spacer sequence that is only integrated on the C57BL/6 allele as part of the reporter gene and has been used before for confirming plasmid and enhancer integration (“integration 1.5 reverse”). For the other two lox sites, forward primers mapping just upstream of and partially overlapping the lox sites (“25/75 forward”) were combined with reverse primers that had been used before for confirming plasmid and enhancer integration and recognise both alleles (“integration 75 reverse”/”lox KI 25 kb initial reverse”). These primer combinations gave rise to PCR products spanning the insert and the surrounding genome. Thus, PCR products having the expected size confirmed the intactness of the inserted sequence and the surrounding genome. Selected PCR products were submitted for Sanger sequencing to confirm further the absence of mutations in the inserted enhancer sequences. The size of the SCR and SCR-rev prevented the amplification of a single PCR product spanning the entire insert. Instead, we performed two PCRs giving rise to partially overlapping PCR products that together span the entire insert and the surrounding genome (“1.5/25/75 forward+SCR-reverse” and “SCR- forward+integration 1.5/25/75 reverse” for SCR, “1.5/25/75 forward+SCR-forward” and “SCR- reverse+integration 1.5/25/75 reverse” for SCR-rev).

To integrate the dual enhancers, we selected individual clonal cell lines from clone 1 before removing the plasmid backbone, i.e., before FlpO treatment. We integrated the second enhancer by transfecting the cells with Cre-recombinase and the second enhancer plasmid. We included both selections to ensure that both enhancers are integrated and selected single clones. We identified positive clones via PCR with the primers described above and removed both backbones by FlpO treatment afterwards. For all cell lines, extensive PCR validation ensured both enhancers were integrated.

### Cas9-targeted sequencing

To verify correct integrations and that the whole locus is not rearranged during the construction of the synthetic locus, we performed targeted nanopore sequencing with Cas9-guided adapter ligation for selected cell lines, both before and after enhancer integration. We designed two small pools of gRNA libraries. The distance between each gRNA is 5kb. Cas9 protein, tracRNA and crRNA were ordered from IDT. We cooperated with the Vienna BioCenter NGS core facility and followed the protocol for Cas9-targeted sequencing using the Cas9 sequencing Kit(SQK-CS9109) from Oxford Nanopore Technologies.

### FACS analysis of mCherry expression

To assess reporter gene activation upon enhancer integration, mCherry levels were measured by FACS. Therefore, 200,000 cells were plated on a 6-well plate in 2i/LIF medium. For each experiment, the two parental reporter cell line clones without enhancer integration and an untransfected v6.5 control were included. After two days, cells were collected by adding trypsin-EDTA (Sigma-Aldrich). Trypsinisation was stopped after incubation at 37 °C for 7 minutes by adding 2i/LIF medium containing 10% serum. Cells were resuspended and centrifuged for 3 minutes at 300 g, the supernatant was removed, and the cell pellet was resuspended in 2i/LIF medium. mCherry-fluorescence was measured with a BD LSRFortessa^TM^ Flow Cytometer (BD Life Sciences - Biosciences).

The resulting FCS files were analysed with the FlowJo^TM^ software (BD Life Sciences – Biosciences, version 10.5.3). We visualised cell populations as histograms normalised to the maximum cell count for each cell line to analyse the fluorescence distribution in a cell population. We calculated the mean and coefficient of variation of mCherry fluorescence with the FlowJoTM software to compare different cell lines and replicates. To account for background fluorescence, we subtracted the mean fluorescence of an untransfected v6.5 control from all values and used the resulting values for the subsequent analysis described below.

We calculated the average and standard deviation of mean fluorescence and plotted the resulting values for each time point in bar graphs (see Figures). To assess statistically significant increases in mCherry fluorescence upon enhancer integration, we performed one-sided paired t-tests on cell lines with enhancer integration compared to their respective parental reporter cell line clone without enhancers.

To compare the distance dependency of different enhancers, we normalised the fluorescence of cell lines with the enhancer at all distances and the parental cell line without enhancer to the respective clone with the enhancer at 1.5 kb. We performed one-sided paired t-tests on the normalised values to identify statistically significant increases compared to the respective parental reporter cell line clone without enhancer.

To compare combinations of enhancers, we calculated the mean mCherry expression of all cell lines, either with one or the combination of two enhancers, and subtracted the mean mCherry expression of clone 1 (no enhancer control) from each mean expression. We calculated the expected additive expression by adding the no enhancer control subtracted values.

### Modelling of expected combined activities

The individual activity of enhancers in the 75kb, 25kb and 1.5kb pads were scaled using the negative control conditions where the three pads contain control sequences (-/-/-). Therefore, they correspond to fold-changes normalised to the basal activity of the core promoter expressed in log2. Hence, for a given pair *P*, log2 additive and multiplicative predicted values were computed using the following formulas:

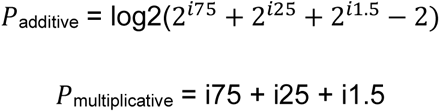

Where *i75, i25 and i1.5* correspond to the log2 individual enhancer activities inserted 75kb, 25kb and 1.5kb upstream of the core promoter, respectively. Finally, we fitted a linear model using log2 activity values and the *lm* function in R:

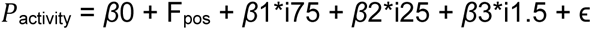

Where *β*0 is the intercept and Fpos corresponds to the position of the two enhancers (+/+/-, +/-/+,-/-/+). *β*1, *β*2 and *β*3 coefficients represent the contribution of each enhancer’s activity and ɛ is the error term. The performance of each model was assessed using R-squared (R^2^) coefficients.

### FACS sorting of mCherry-populations

Upon integration of the *Nanog* enhancer at 1.5 kb, we observed some clones with a fraction of mCherry-negative cells. We sorted mCherry-positive and −negative cells with a BD FACSAria™ III Cell Sorter (BD Life Sciences - Biosciences). Sorting gates were determined by measuring mCherry fluorescence of a clone without a fraction of mCherry-negative cells: All cells with mCherry signal lower than the lowest-expressing cells of that clone were sorted as negative, and all remaining cells were sorted as positive. Sorted cells were kept in culture, and mCherry levels were analysed with a BD LSRFortessa^TM^ Flow Cytometer (BD Life Sciences - Biosciences) every few passages.

### Luciferase assays

For luciferase assays, we used a pGL3 plasmid with the Firefly luciferase coding sequence followed by a poly-adenylation signal under the control of the SV40 promoter (Promega). The same enhancer sequences we integrated into the reporter locus were amplified by PCR from *castaneus* mouse strain DNA and inserted downstream of the poly-adenylation signal by Gibson assembly. The primer sequences used to generate these plasmids are indicated in Supplementary Table 3. Luciferase assays were slightly adapted but otherwise performed as described previously (Thomas et al. 2021). 10,000 cells from the v6.5 cell line were plated on a 96-well plate and immediately transfected with 120 ng of enhancer-luciferase plasmid, 4 ng of Renilla control plasmid (Promega) and 0.62 μL of Lipofectamine 2000 Transfection Reagent (Invitrogen^TM^). The medium was removed 5-7 h after transfection, and fresh 2i/LIF medium was added. Firefly and Renilla luciferase activity were measured 48 h after transfection. Background-subtracted Firefly measurements were normalised to background-subtracted Renilla values. The resulting values were normalised to the control plasmid without enhancer integration. To detect statistically significant increases in luciferase activity compared to the no-enhancer control, we performed one-sided Welch one-sample t-tests on the resulting normalised values to assess whether they are significantly higher than 1 (the value of the no-enhancer control).

### Enhancer KO and RT-qPCR

Enhancers at the Map4k3 locus were deleted in R1 WT ESCs as described above. RNA was extracted from confluent 6-well plates, and resulting Map4k3 expression levels were measured by RT-qPCR as described before^23^. Expression levels of each replicate were normalised first to Rpl13a and then to WT. To assess statistically significant decreases in expression upon enhancer deletion, we performed one-sided Welch one-sample t-tests on the resulting WT-normalized values to assess whether they are significantly lower than 1 (as all values are normalised to WT, a value of 1 corresponds to WT expression levels).

## Additional Resources

### Supplemental Figure legends

**Supplemental Figure 1:**
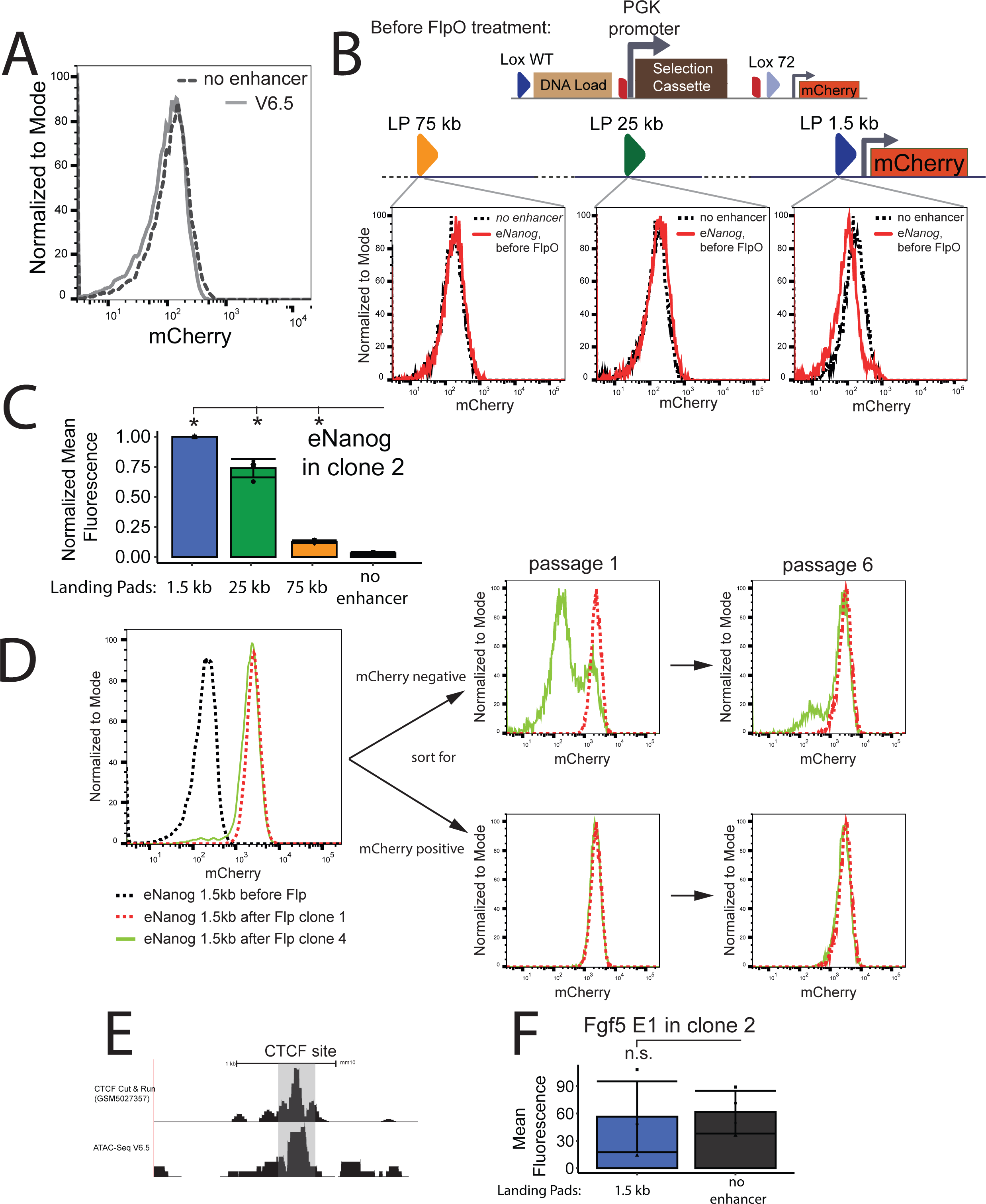
Related to Figure 1. A: FACS plot showing a selected example of clone 2 no enhancer and the parental cell line v6.5 B: Top: schematic of the synthetic locus organisation before removal of the backbone plasmid, Bottom: representative FACS plots of different *eNanog* integrated clones at the indicated LPs before removal of the plasmid backbone C: Quantification of mean mCherry expression of *eNanog* integrated at the indicated landing pads in clone 2. All mean expressions were normalised to the 1.5 kb integration. Statistically significant different signals (p < 0.05, one-sided paired t-tests) are marked by stars. D: FACS plots of mCherry expression in selected clones after backbone plasmid removal; mCherry negative (top) and positive (bottom) cells were sorted and analysed after 1 (left) or 6 (right) additional passages E: UCSC browser track surrounding potential CTCF sites and ATAC-seq signal F: Representative FACS plots showing an individual clonal cell line with the Fgf5 E1 enhancer integrated at the 1.5 kb LP in clone 2.

**Supplemental Figure 2:**
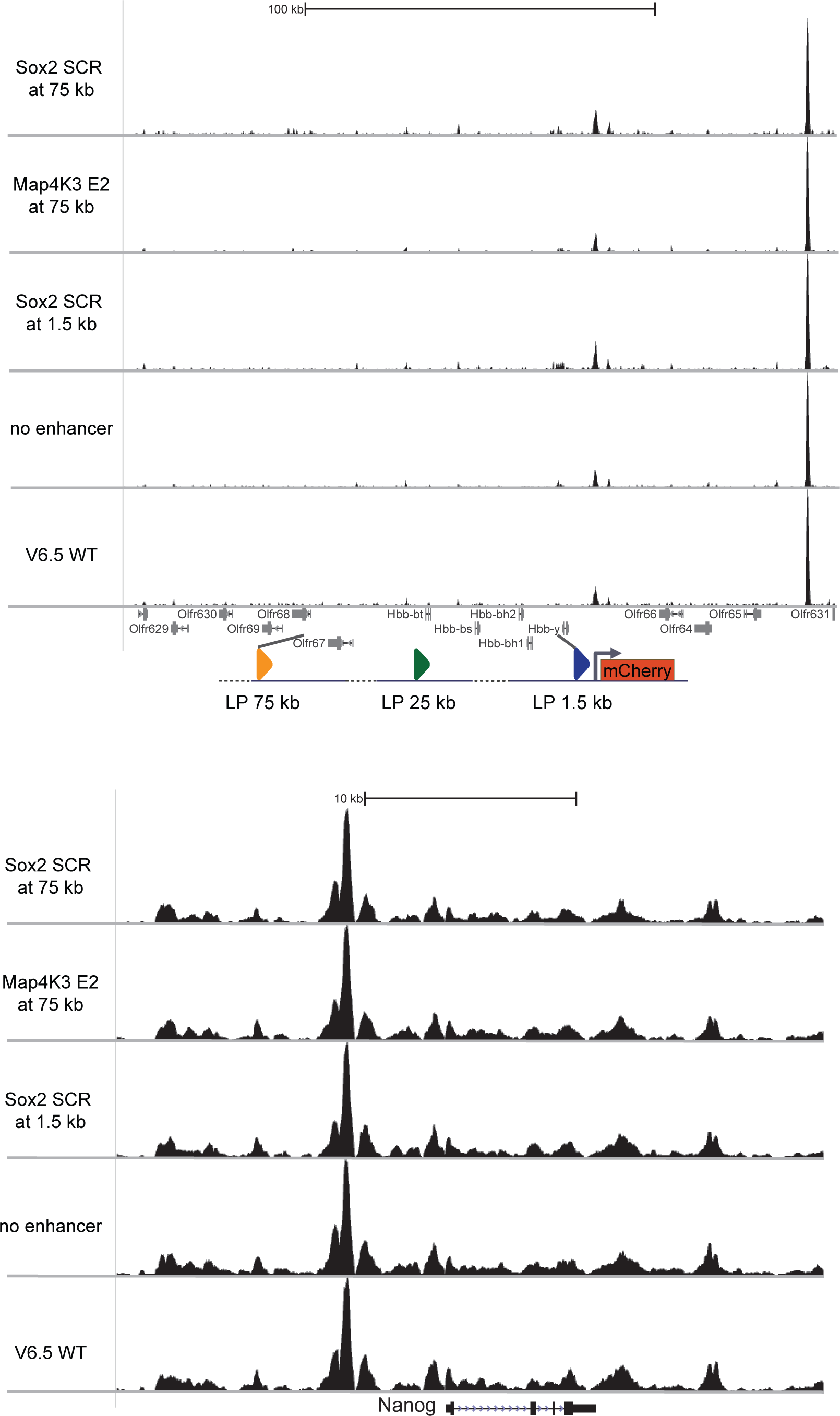
Related to Figure 1. UCSC browser track showing ATAC seq signal at the β-globin locus (top) and the *Nanog* locus (bottom) of the parental cell lines v6.5, the no enhancer control and different enhancers integrated at different LPs.

**Supplemental Figure 3:**
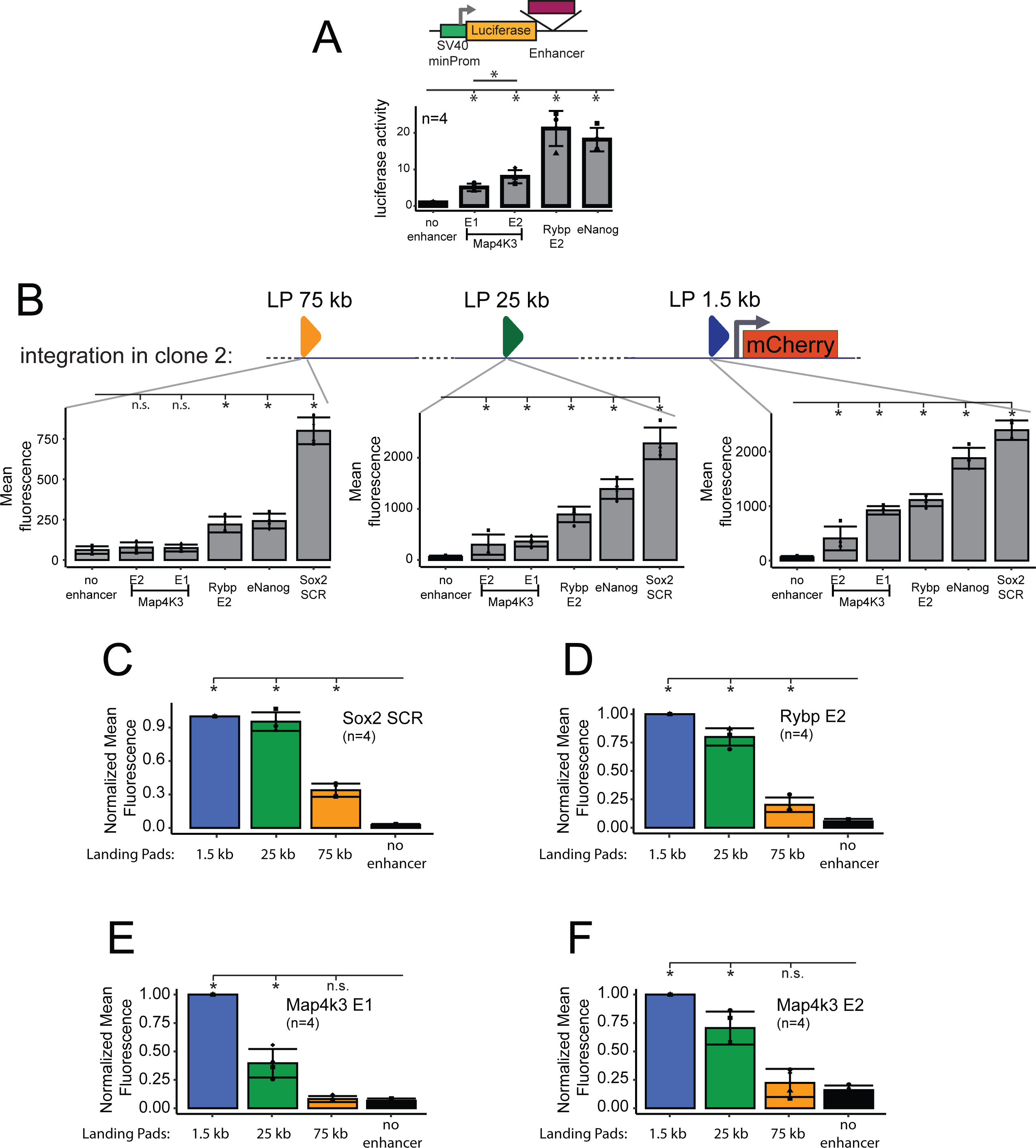
related to Figure 2. A: Luciferase Assay of the selected short enhancers. Top: schematic depicting the integration of the enhancers. Statistical significance was tested by one-sided Welch one-sample t-tests on the resulting normalised values to assess whether they were significantly higher than 1 (the value of the no-enhancer control). B: Quantification of mCherry expression of the five indicated enhancers compared to the no enhancer control at three different LPs. Left: integration at 75 kb, middle: integration at 25 kb, Right: integration at 1.5 kb. Statistically significant signals are marked by stars compared to the no enhancer control (p < 0.05, one-sided paired t-tests). n.s.: not significant, p>0.05. C-F: Quantifying mean mCherry expression of indicated enhancers integrated at the indicated landing pads. All mean expressions were normalised to the 1.5 kb integration. Statistically significant different signals (p < 0.05, one-sided paired t-tests) are marked by stars.

**Supplemental Table 1:**
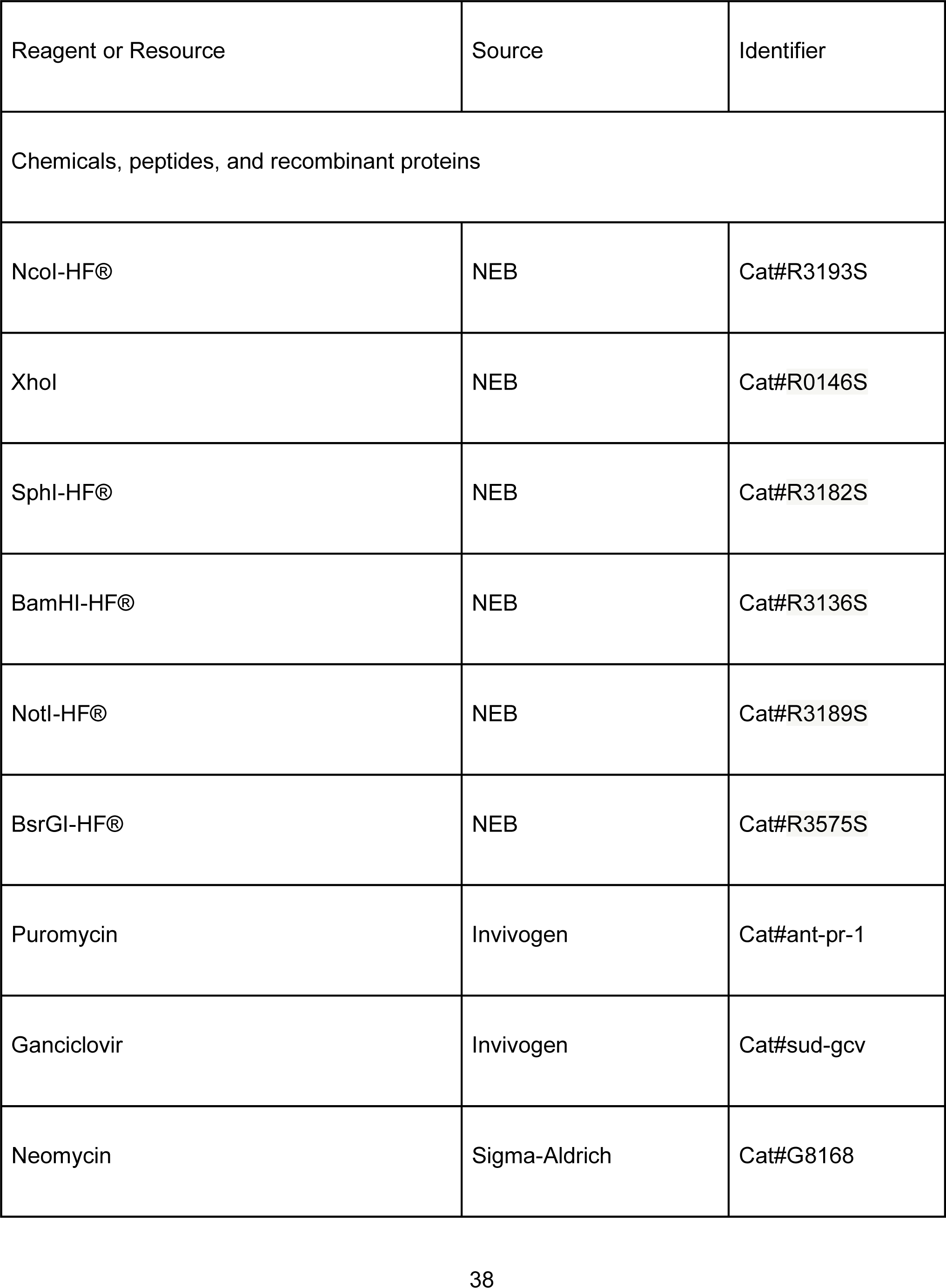
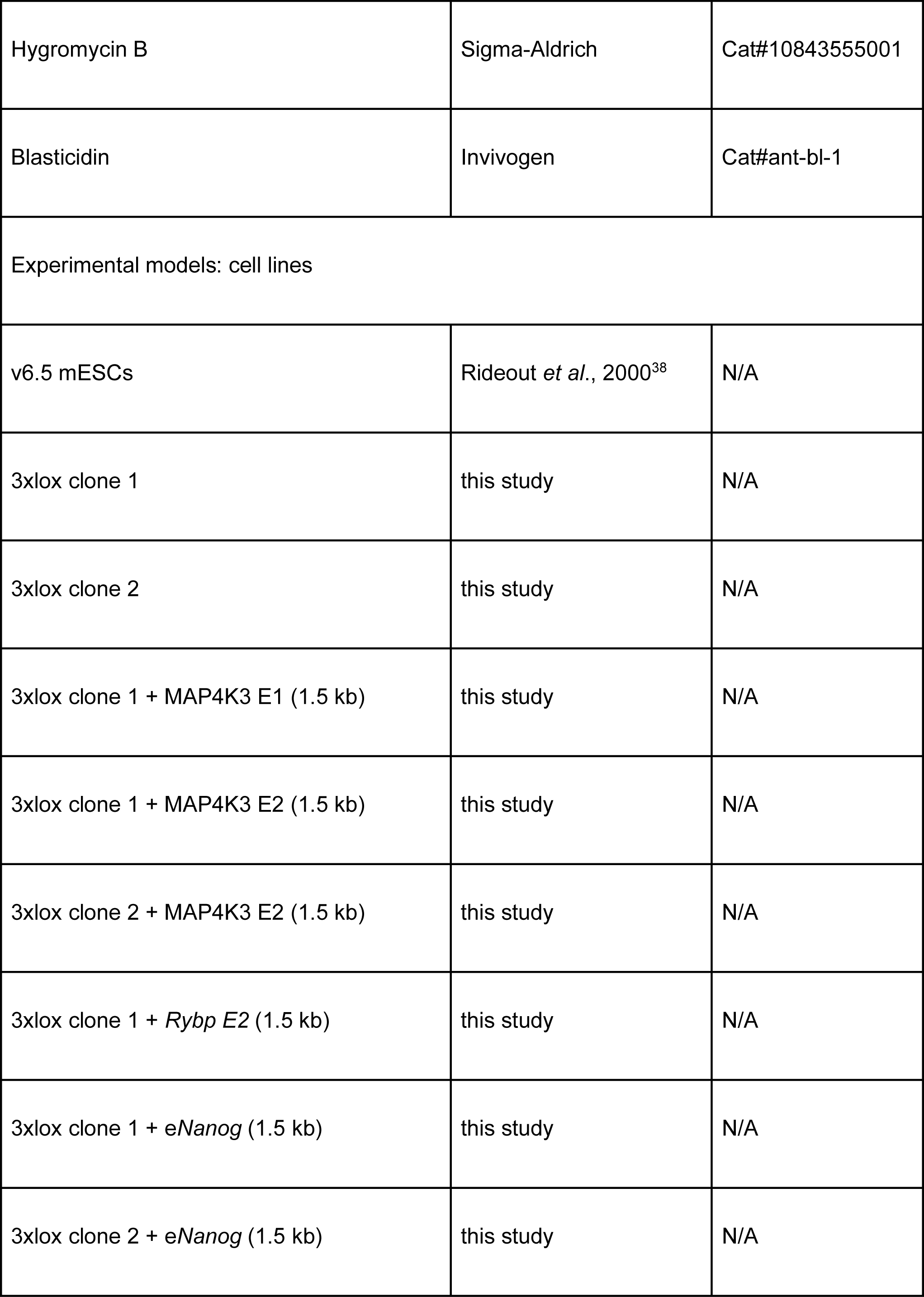
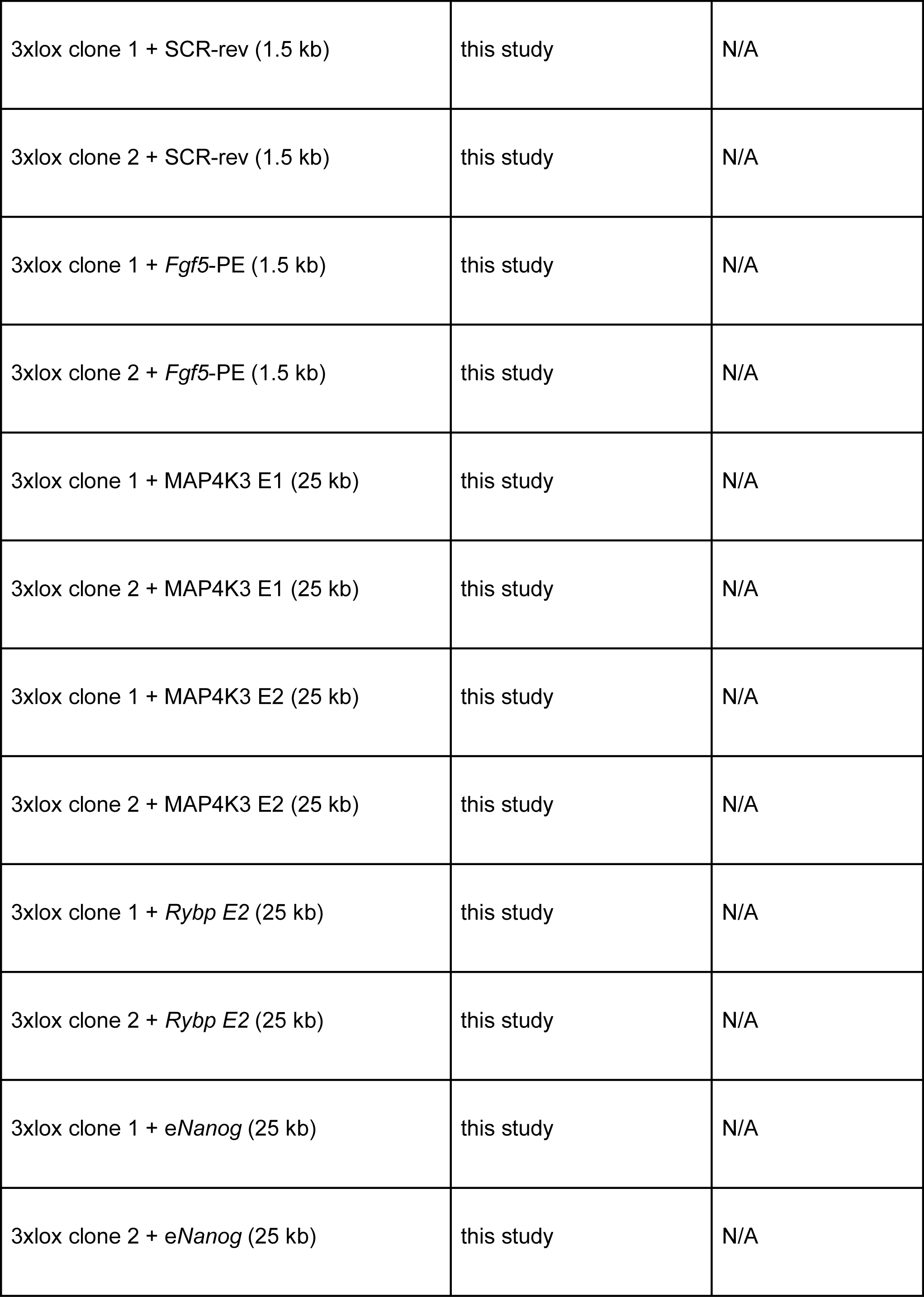
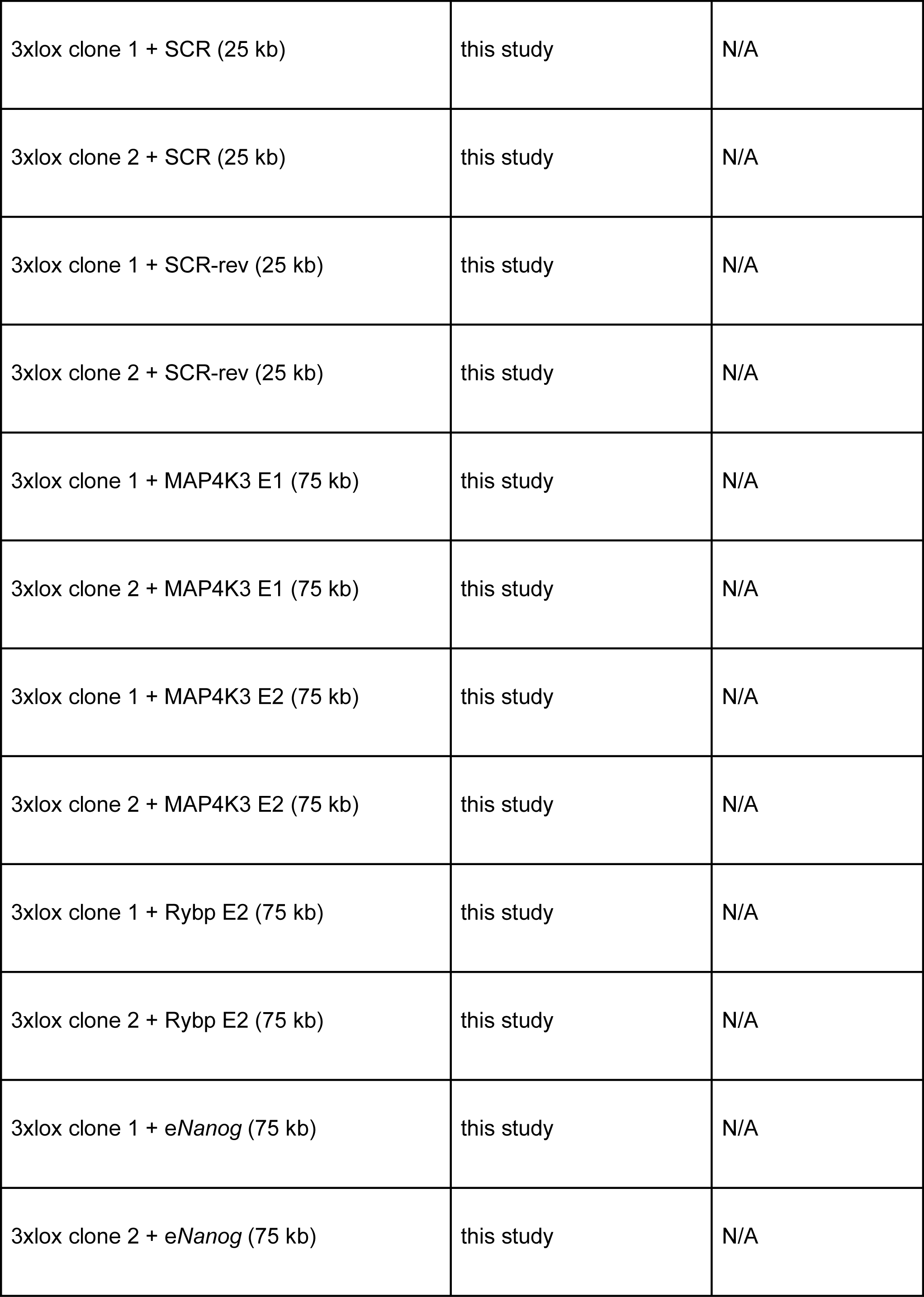
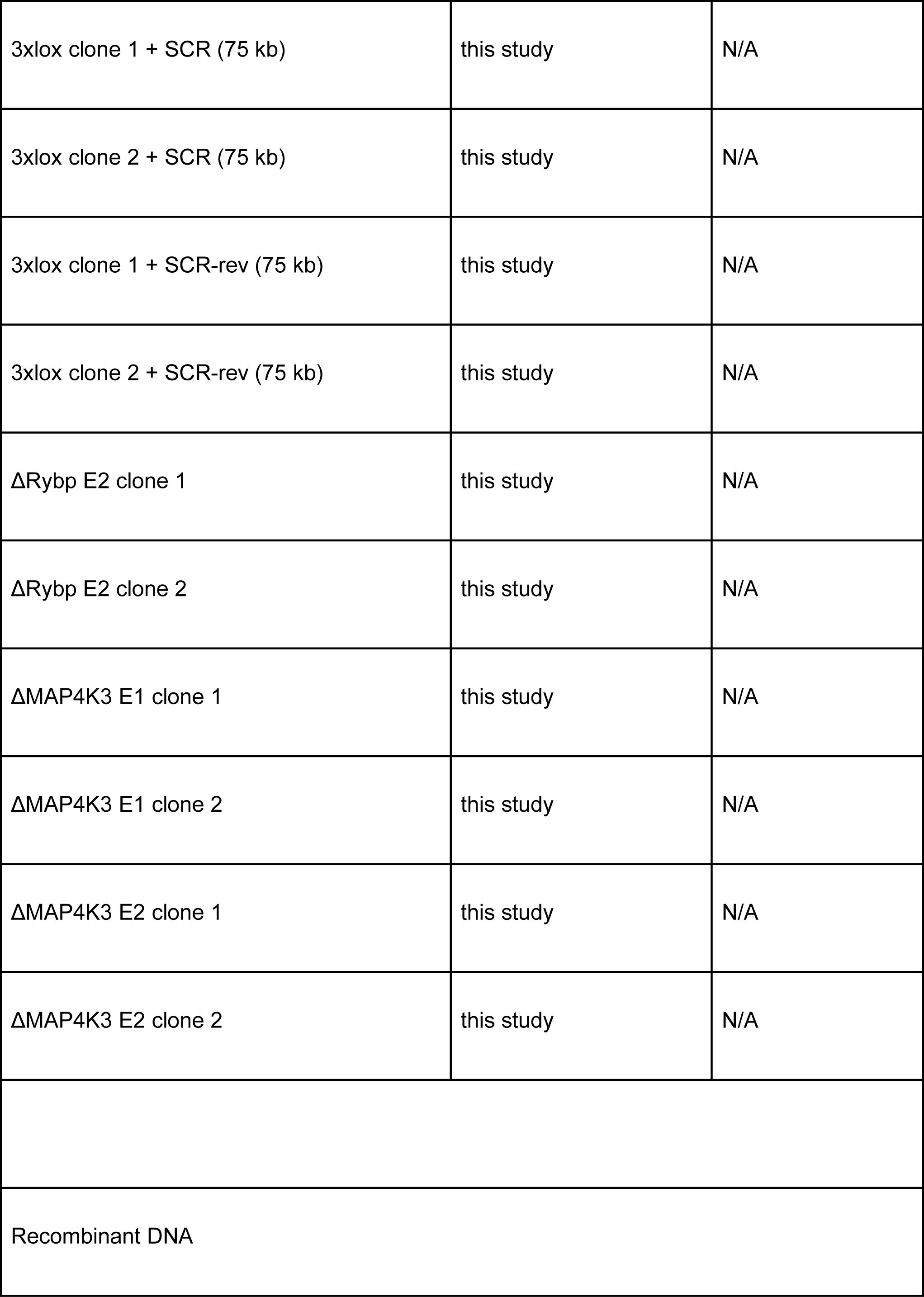
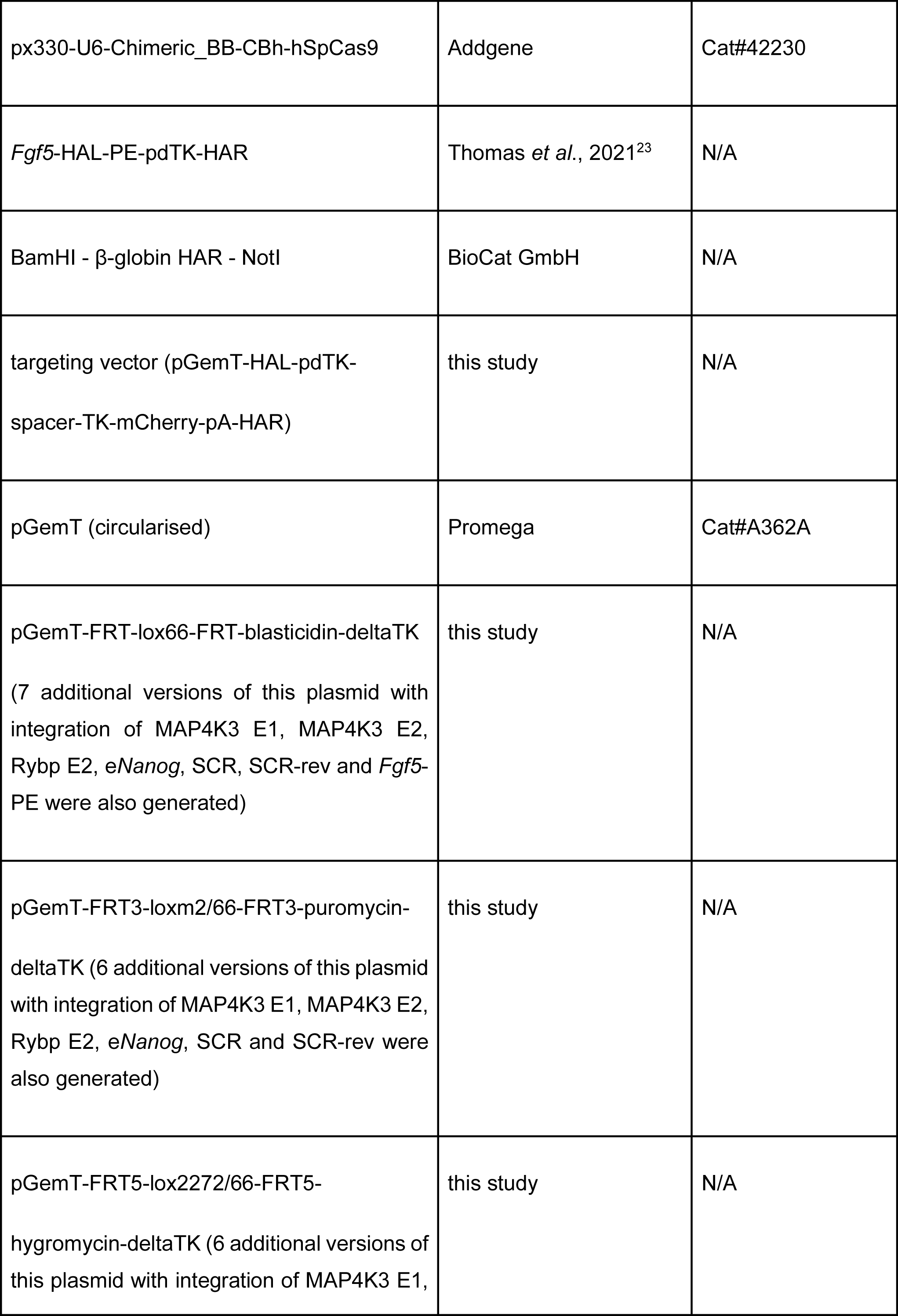
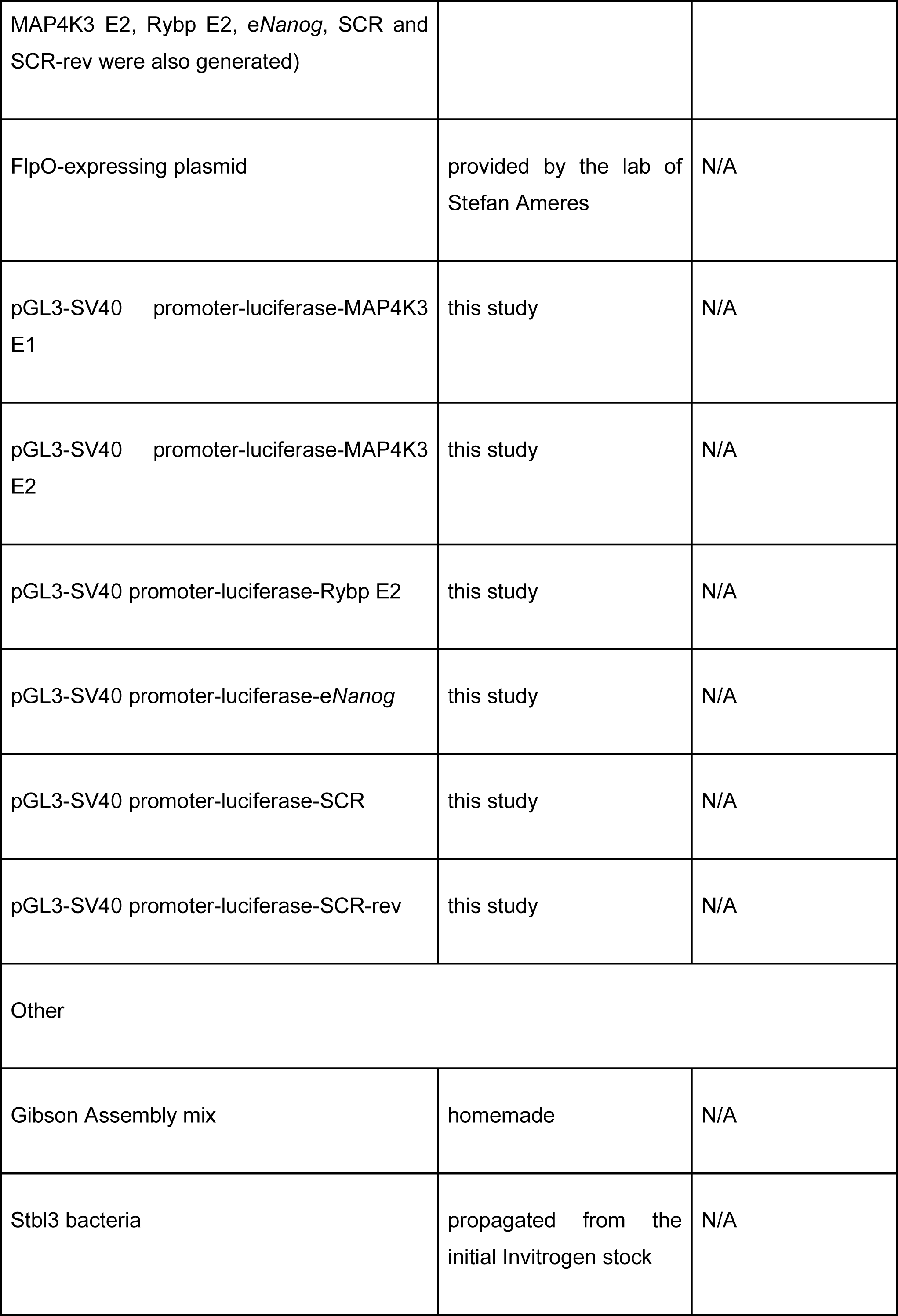
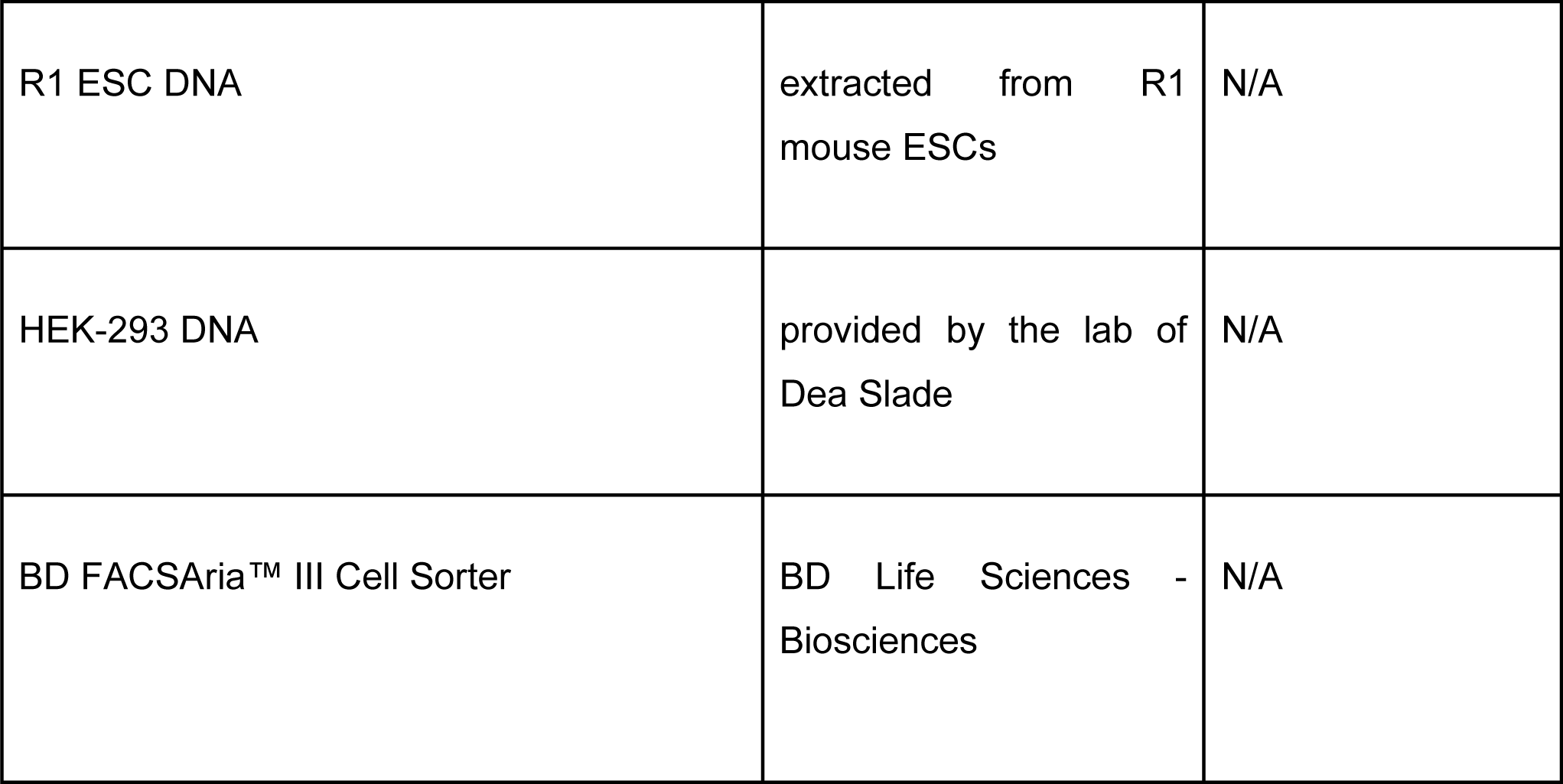
Key Resource Table.

**Supplemental Table 2:**
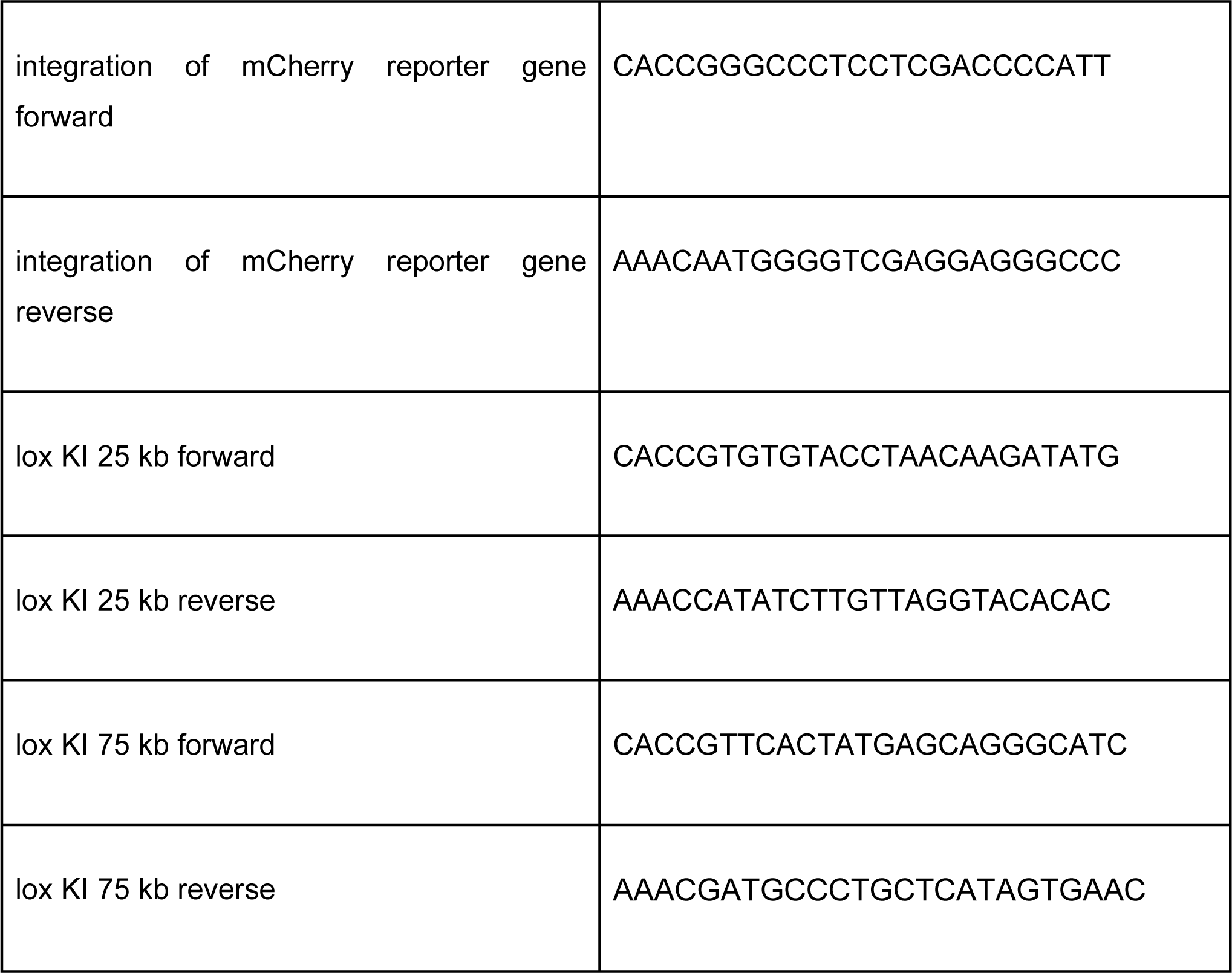
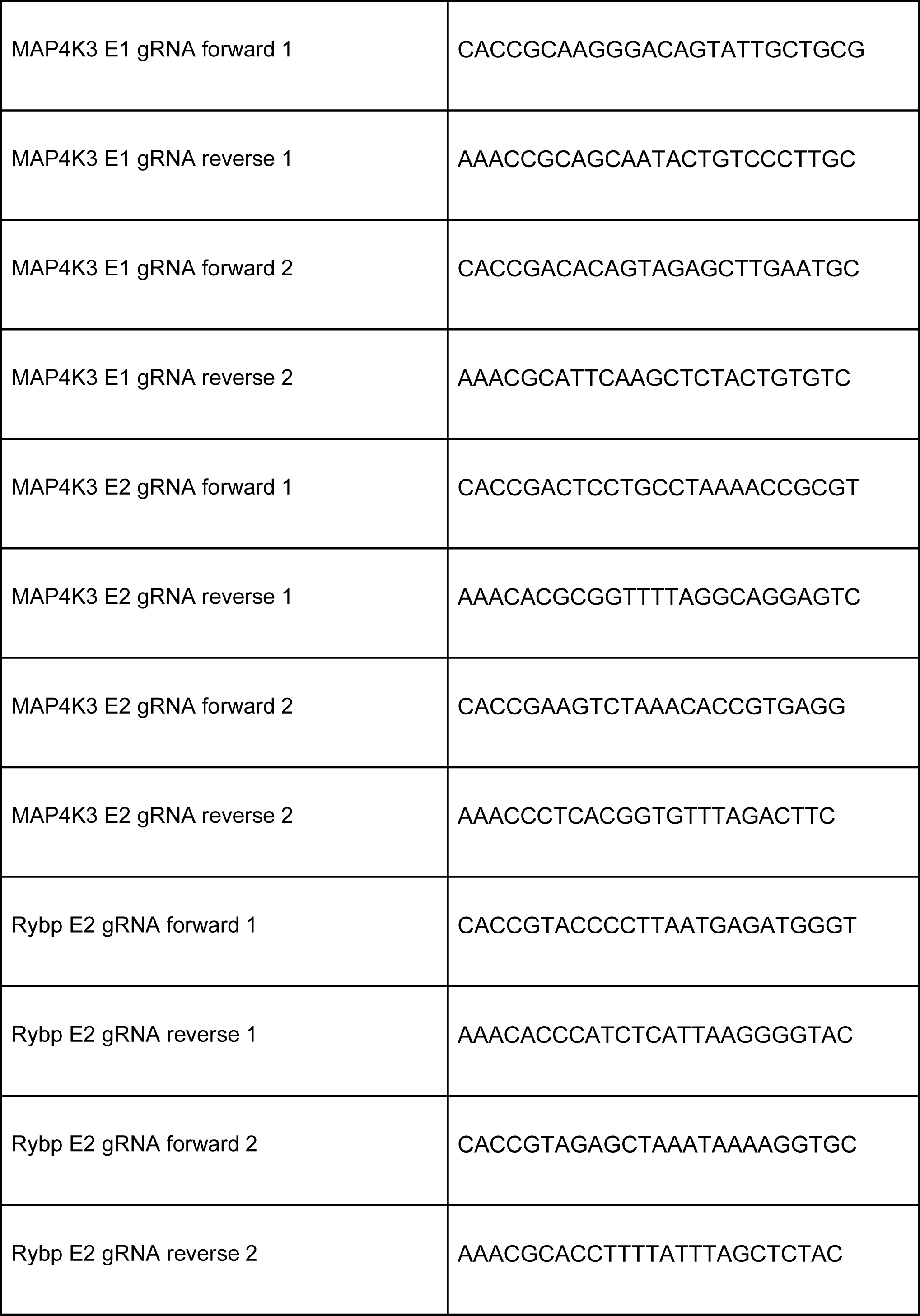
gRNA sequences.

**Supplemental Table3:**
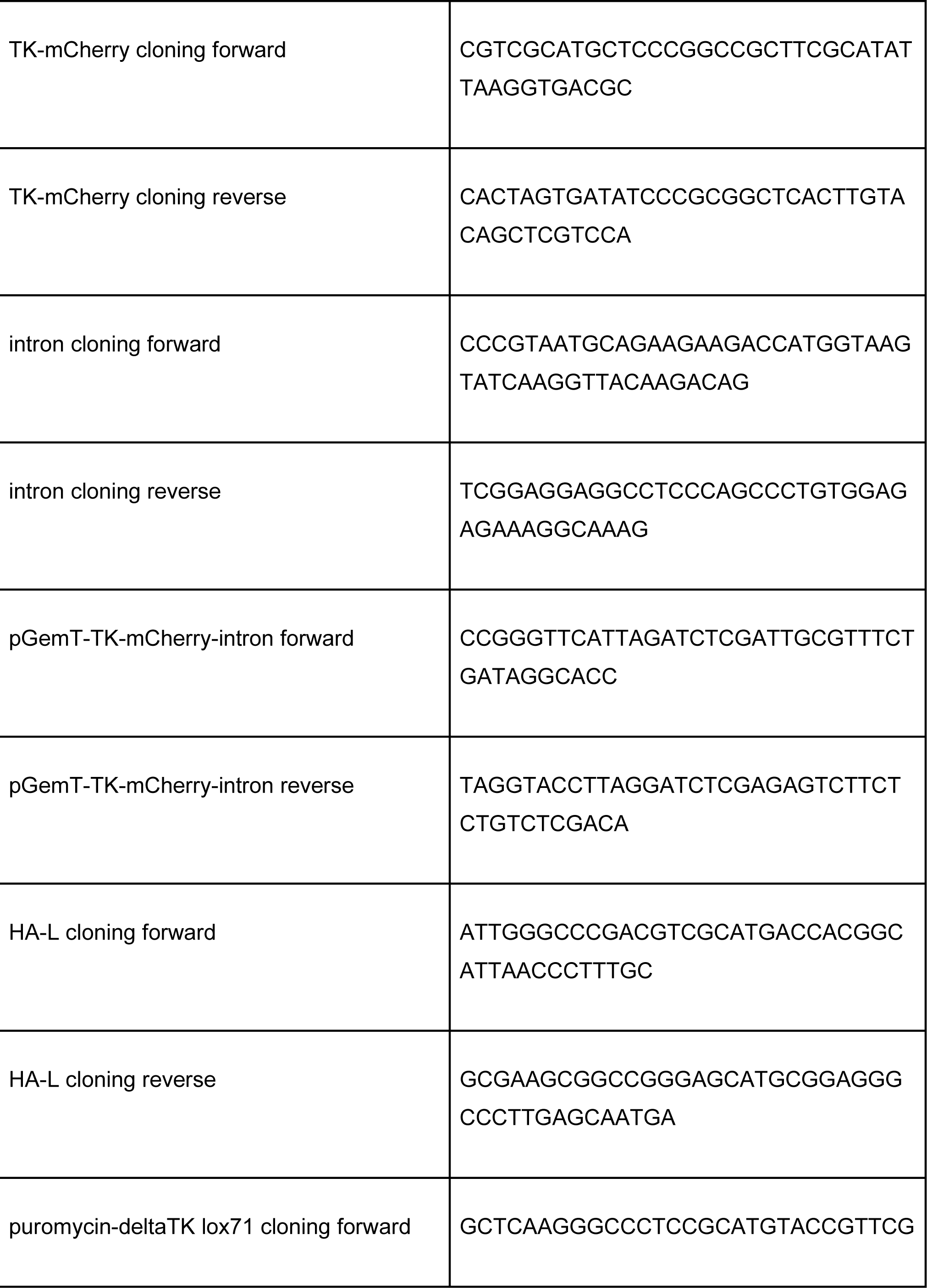
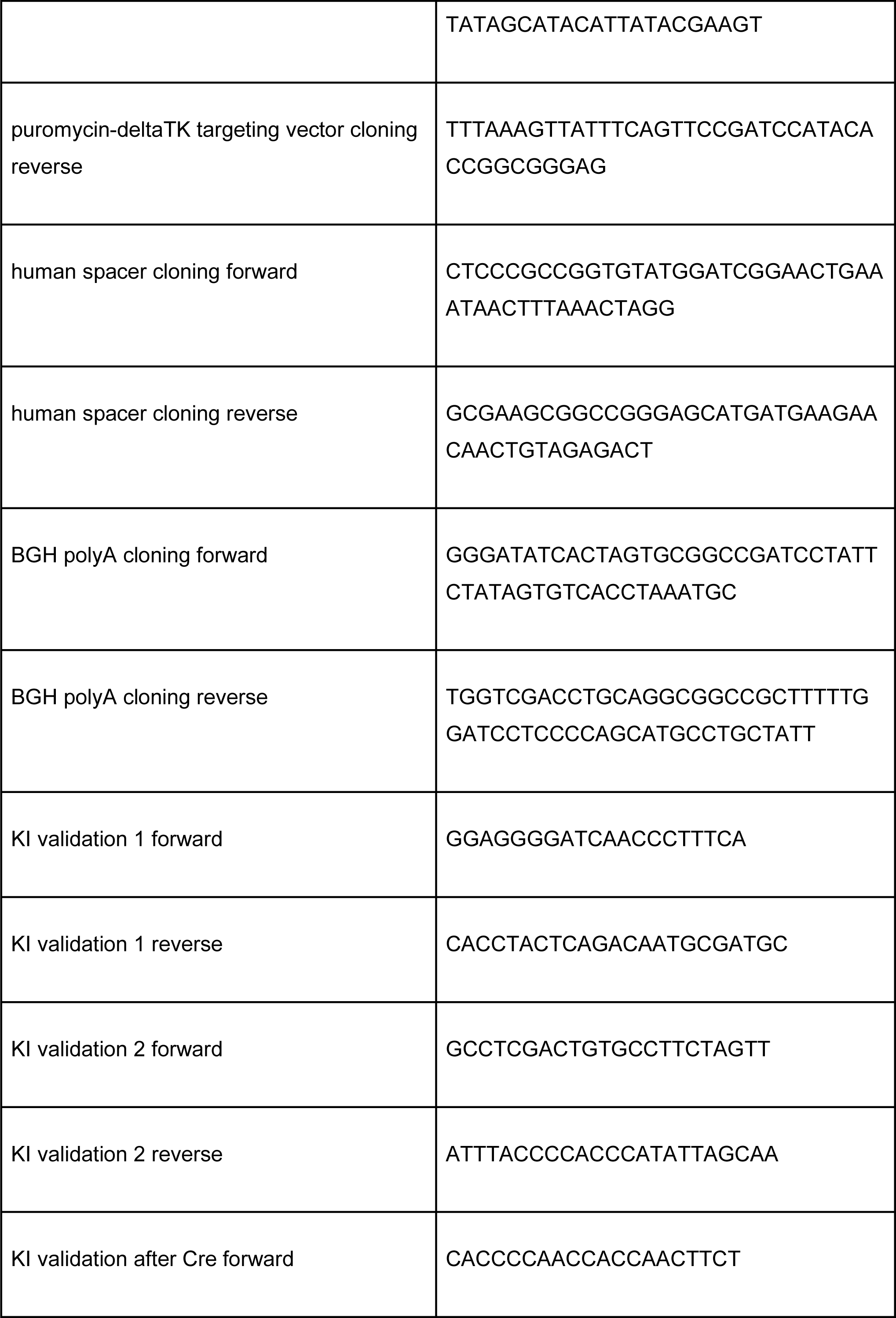
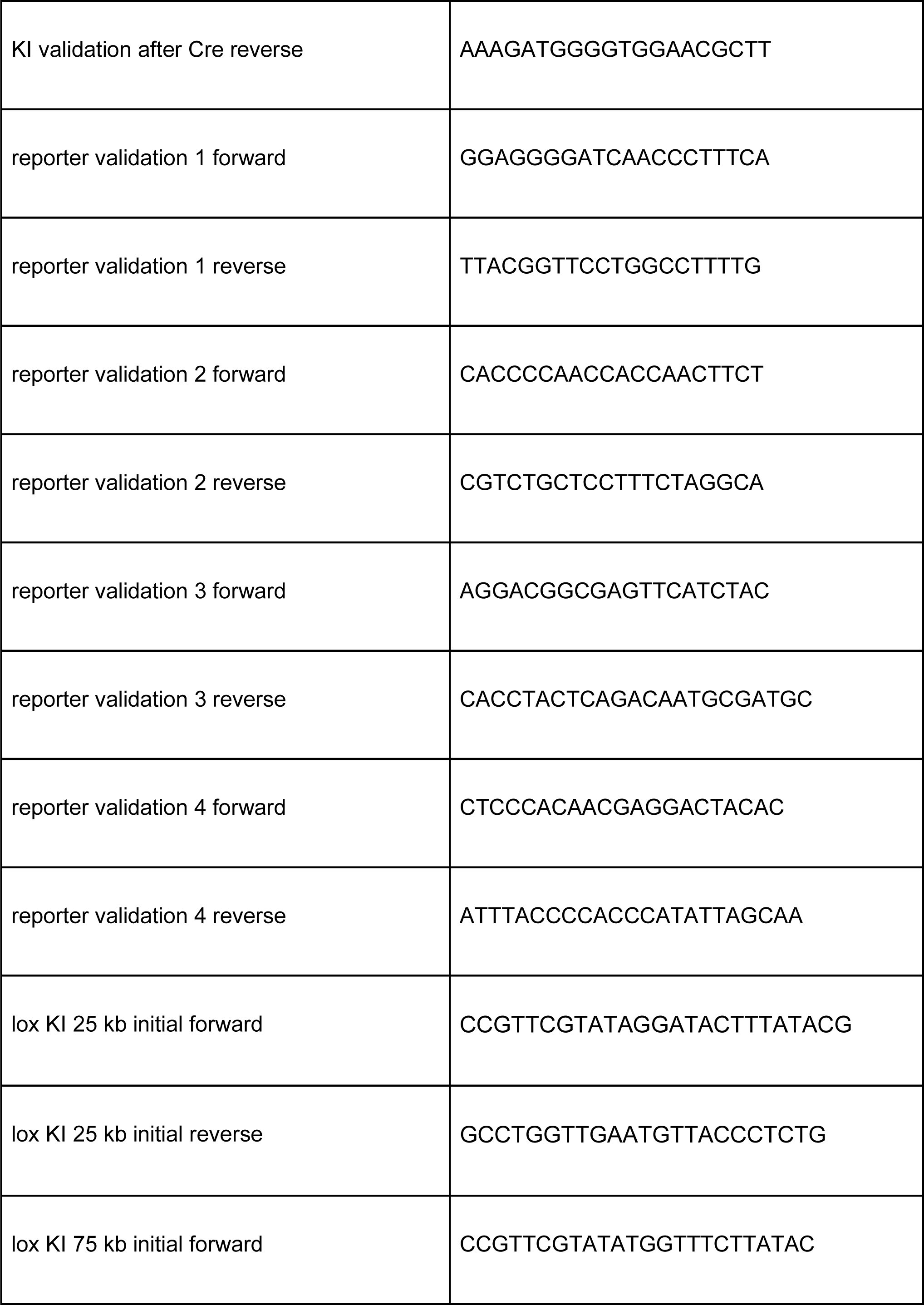
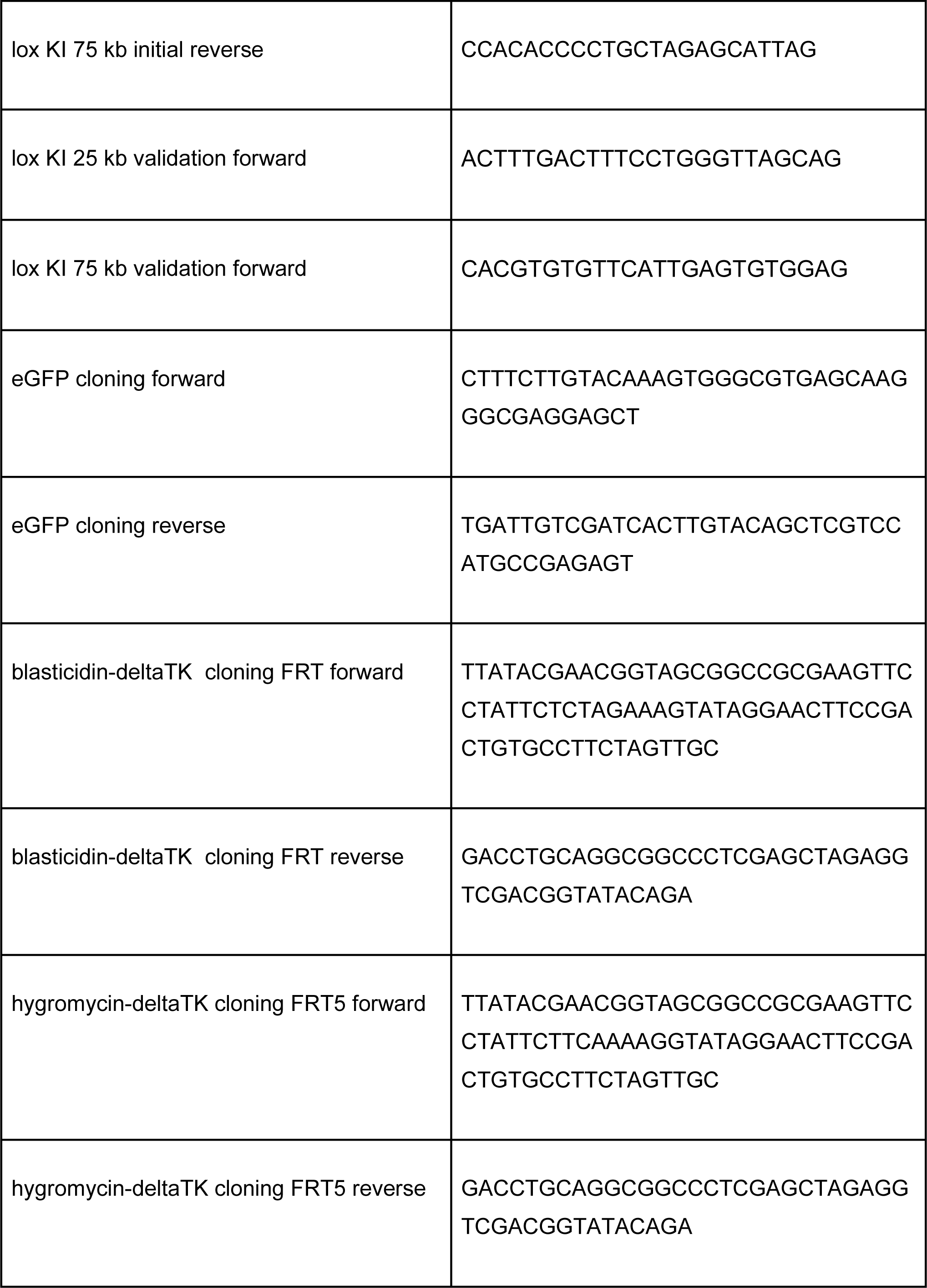
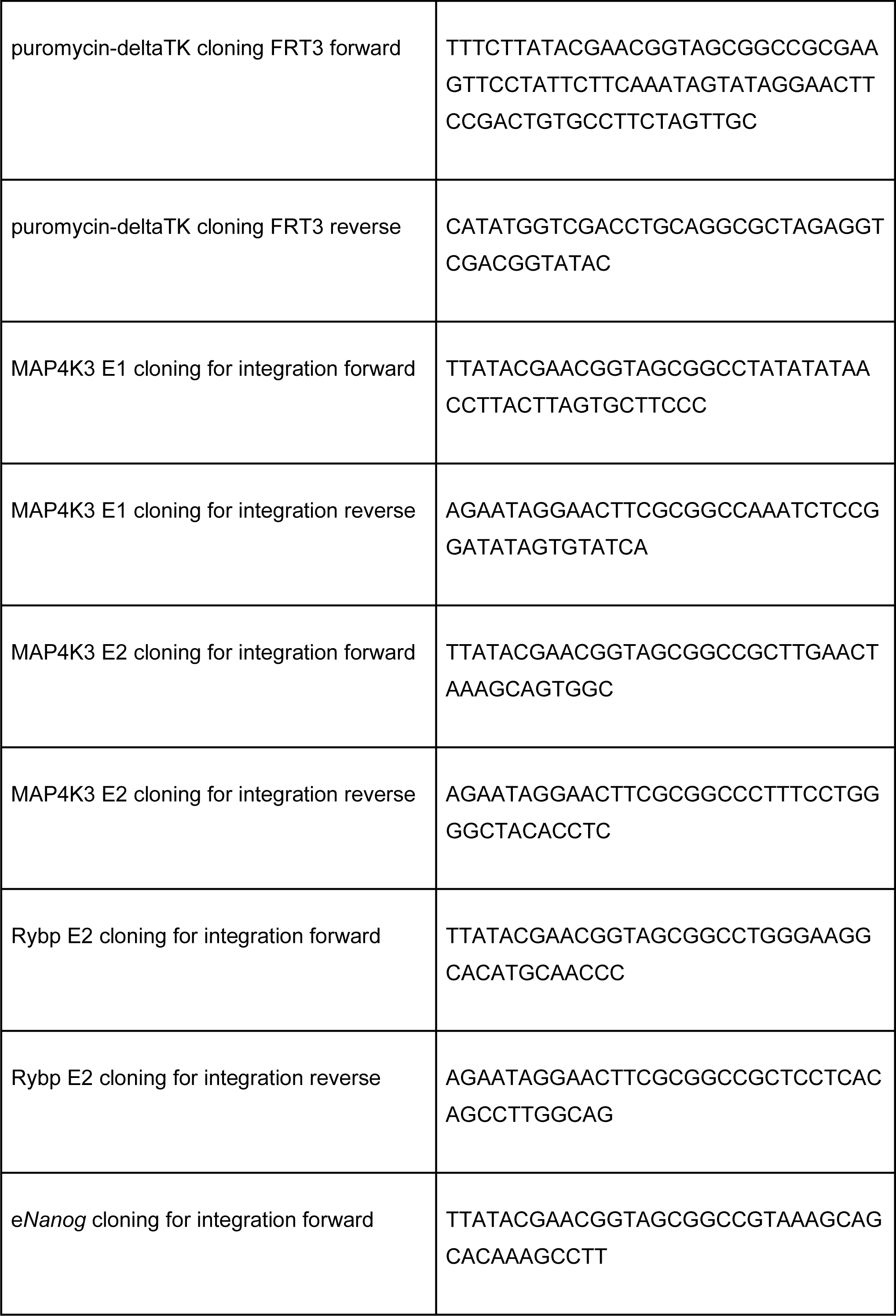
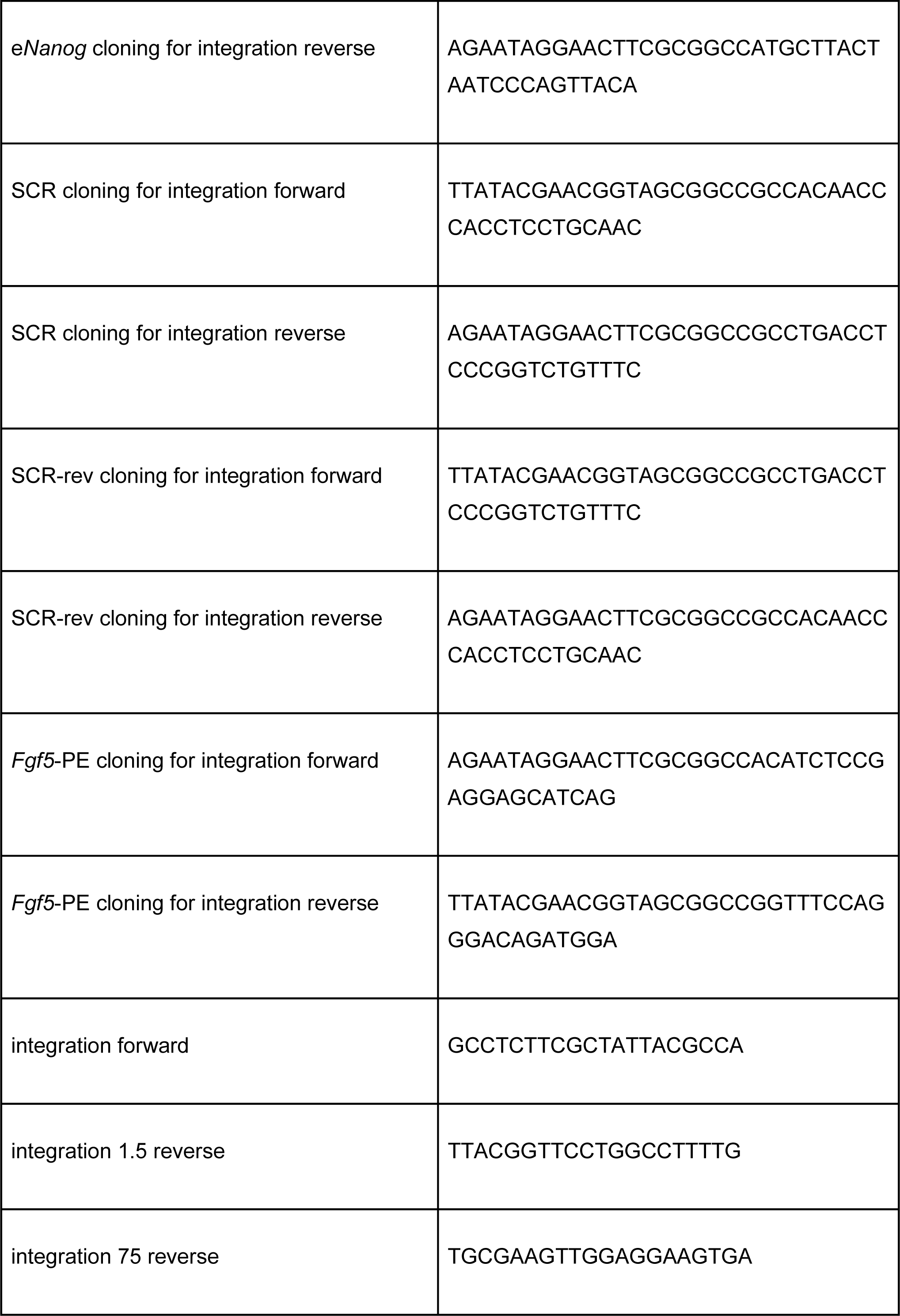
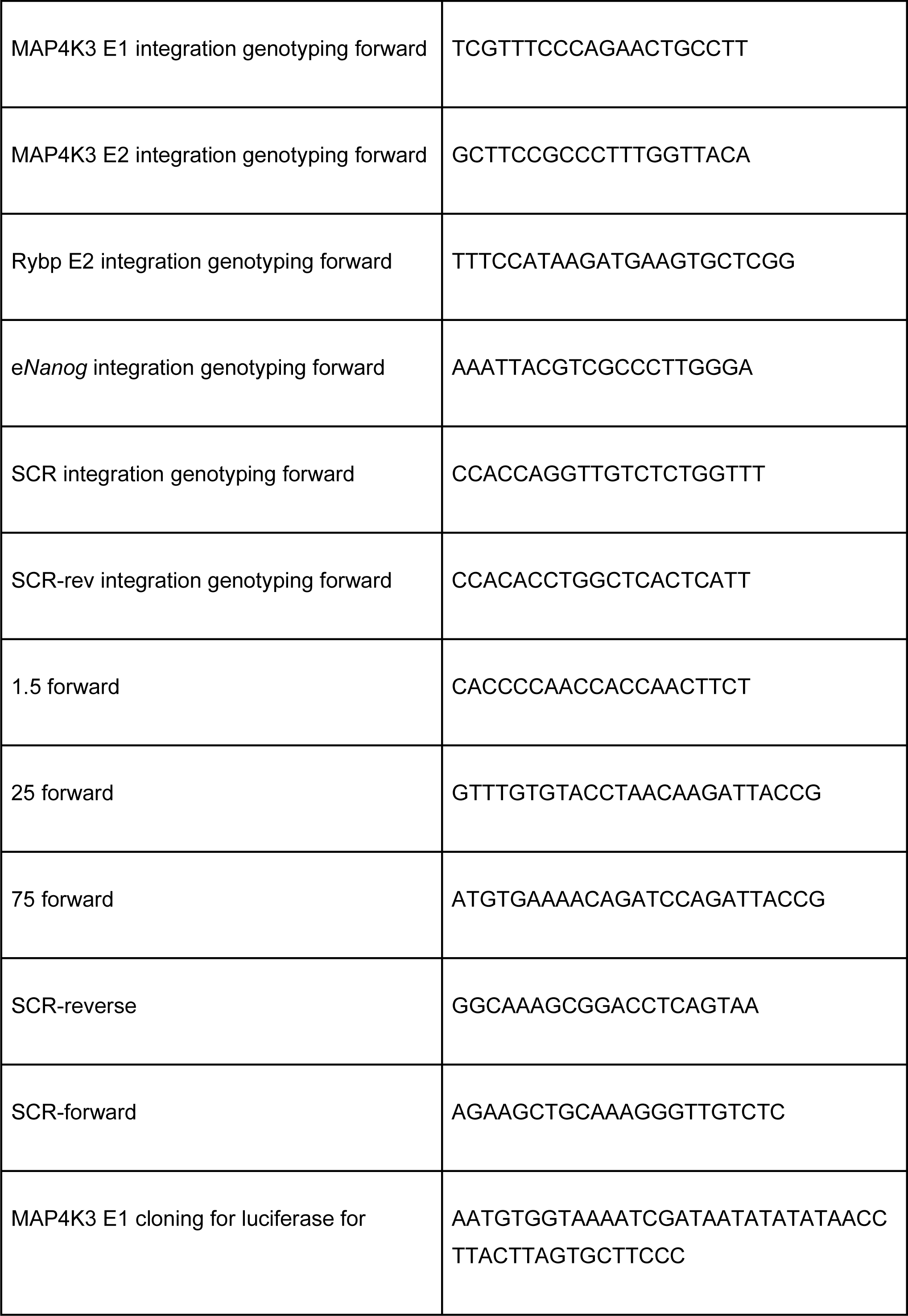
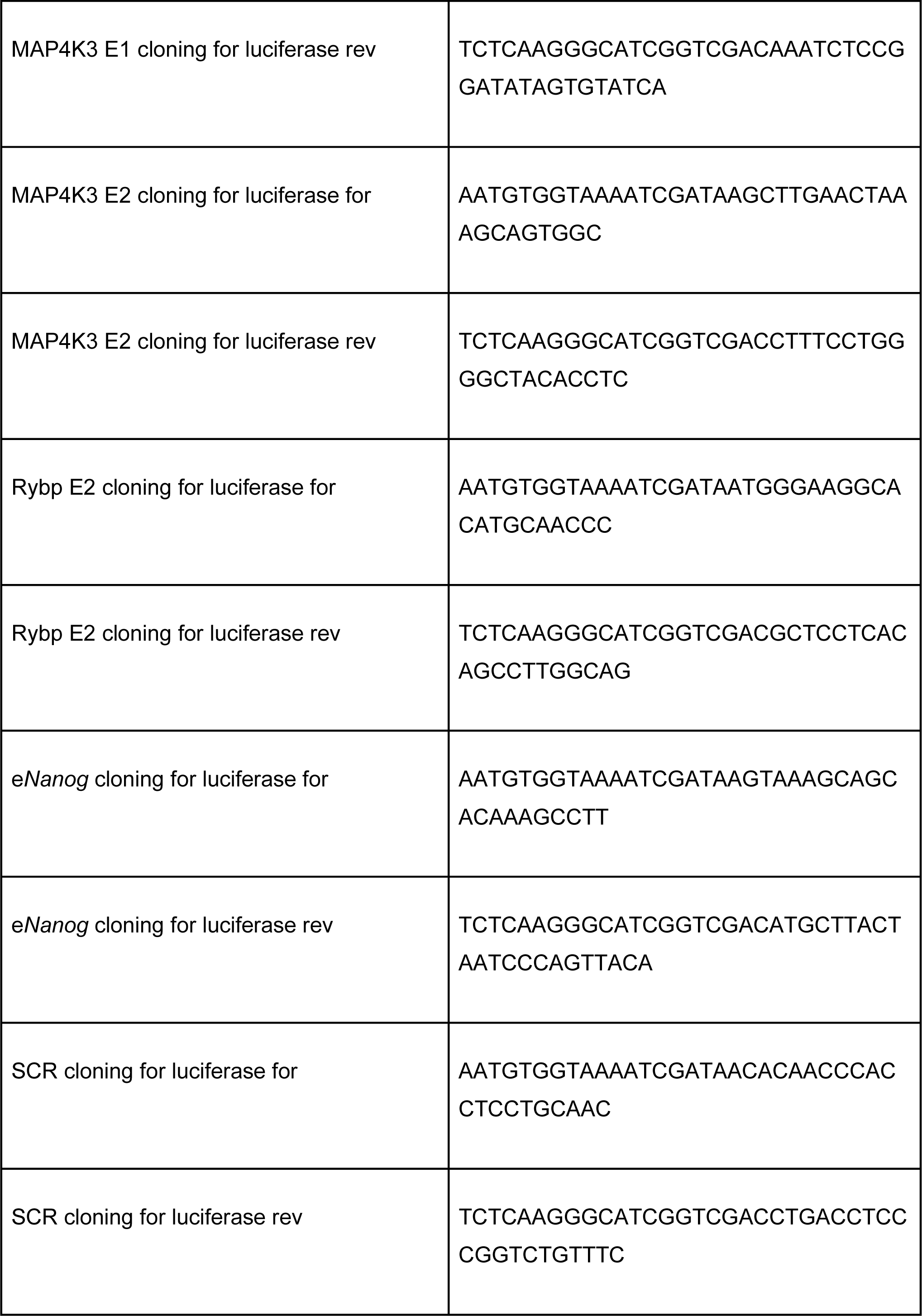
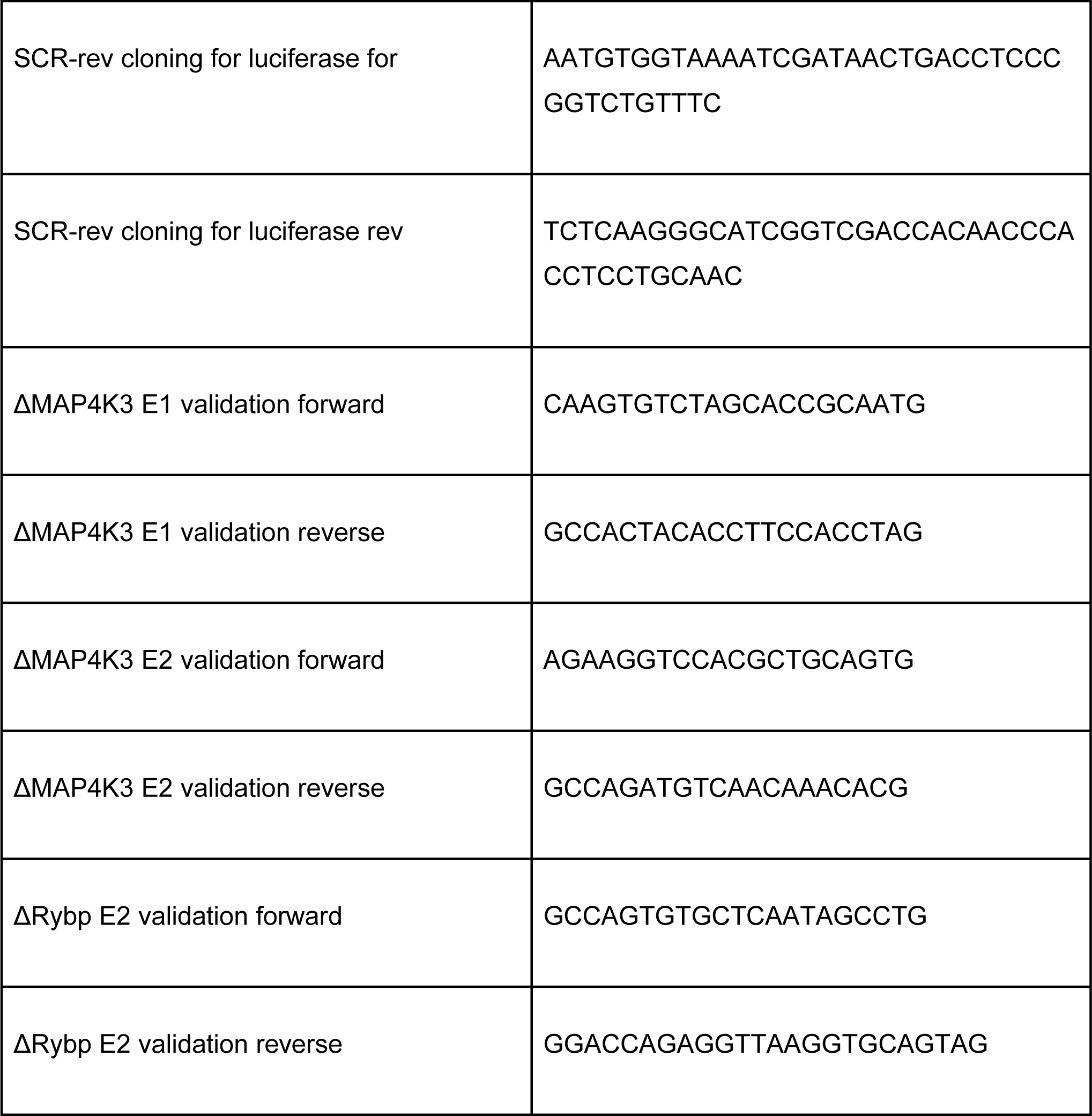
PCR primers.

**Supplemental Table 4:**
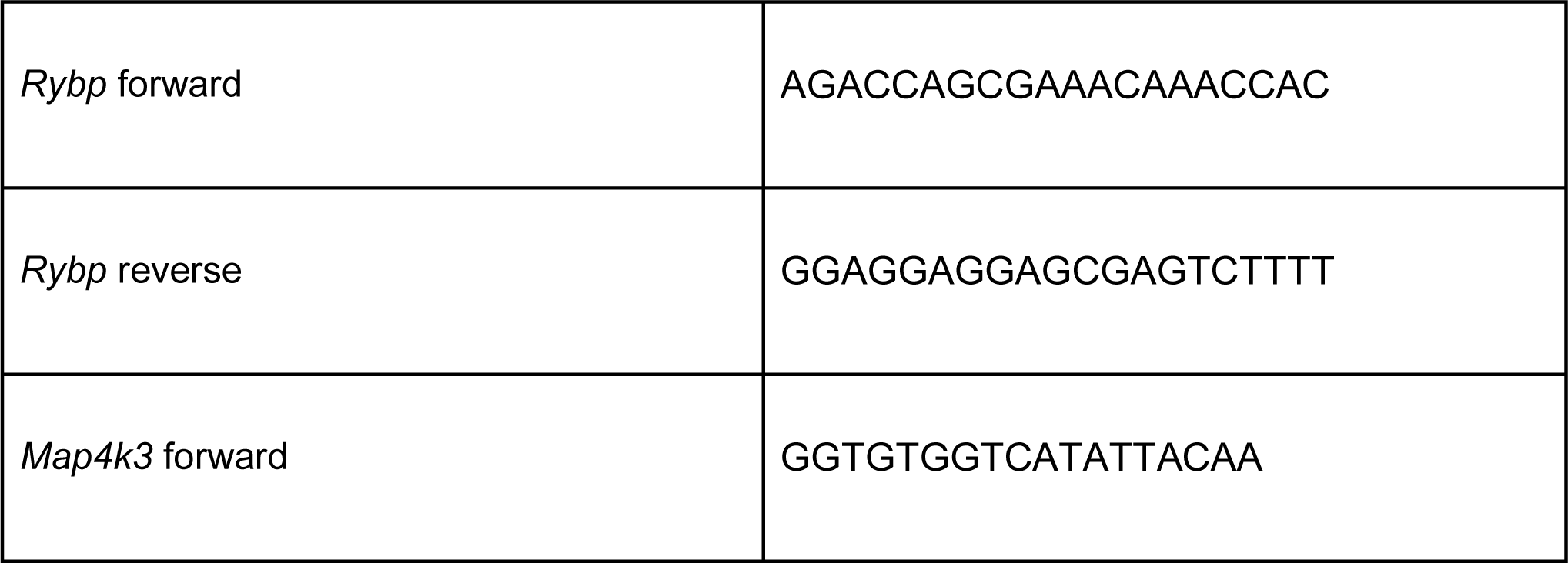
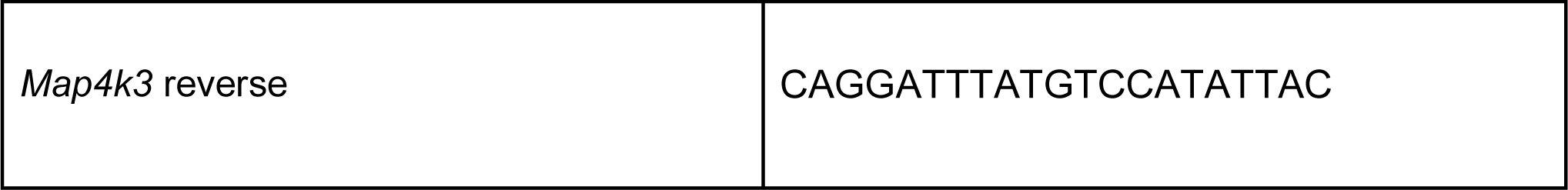
RT-qPCR primers.

**Supplemental Table 5:**
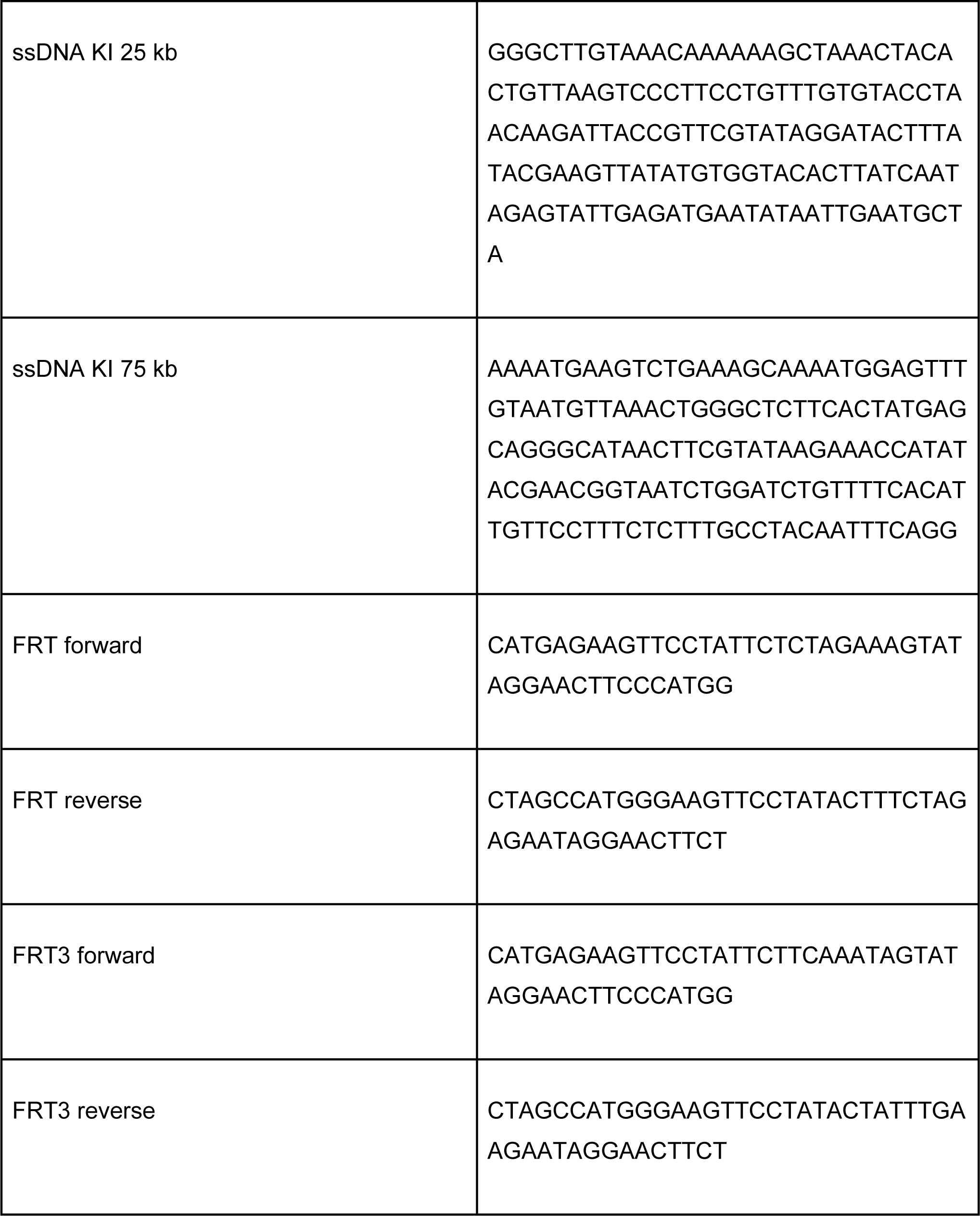
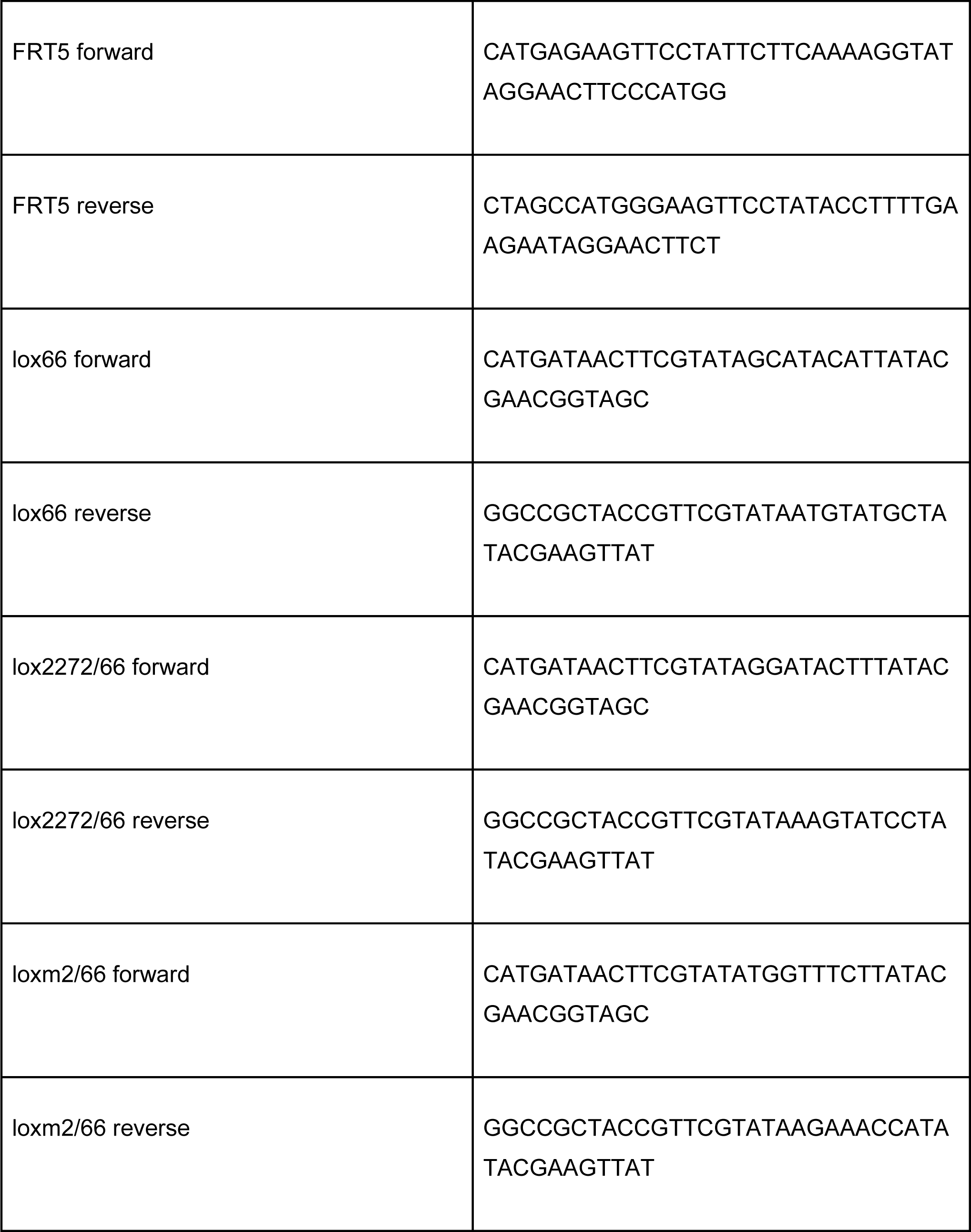
DNA oligos.

**Supplemental Table 6:** Summary of Landing Pads and integration efficiencies.

|  | **lox71 /**<br>**lox66** | **FRT** | **Selection Cassette** | **percentage of**<br>**positive clones** |
| --- | --- | --- | --- | --- |
| **Landing Pad 1.5 kb** | loxP | FRT | Blasticidine-deltaTK | 25% |
| **Landing Pad 25 kb** | Lox2272 | FRT5 | Hygromycin-delta TK | 19% |
| **Landing Pad 75 kb** | loxm2 | FRT3 | Puromycin-delta TK | 24.30% |

**Supplemental Table 7:**
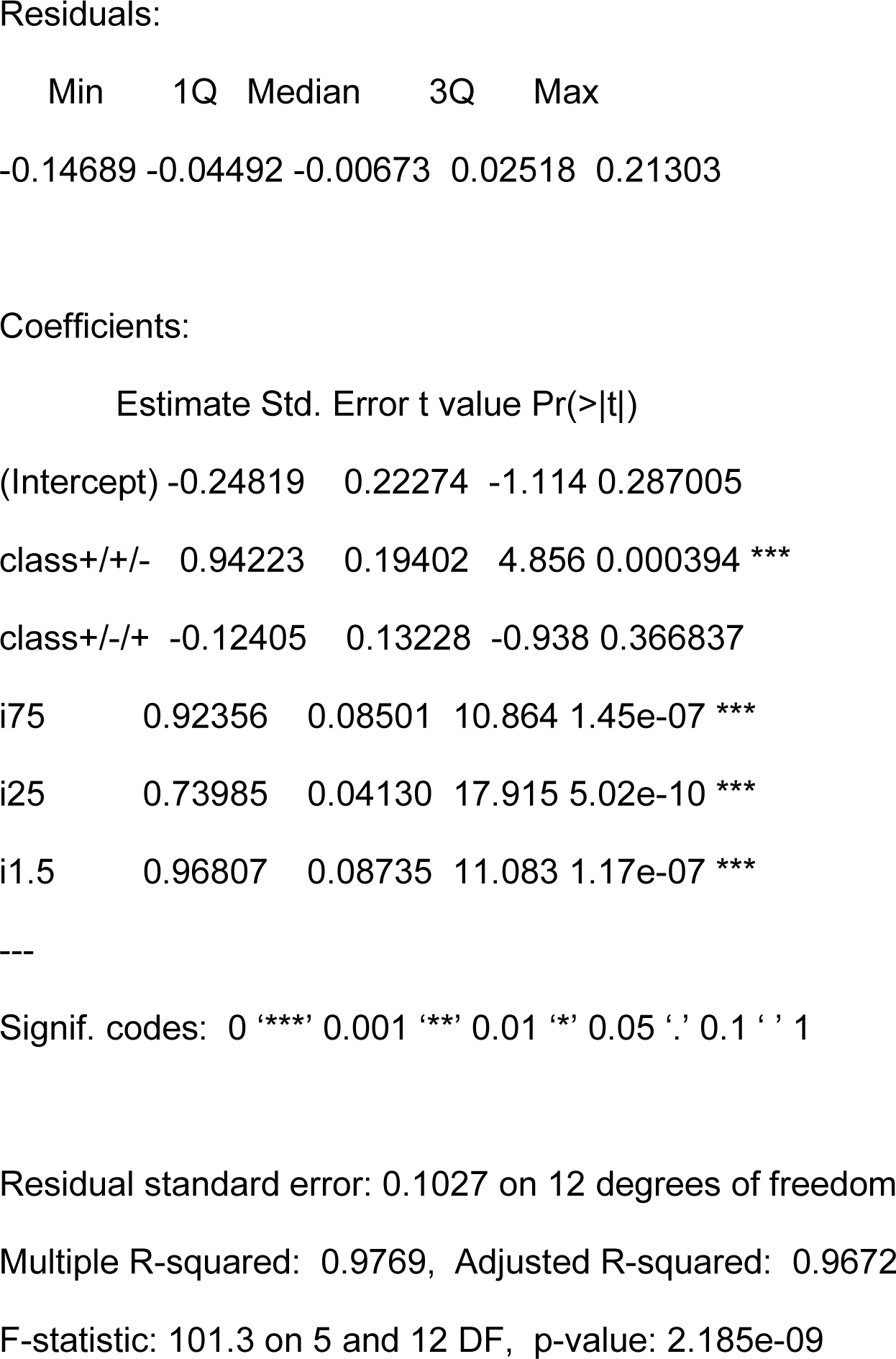
Results from the Linear Modelling.

## Notes

### Competing Interest Statement

The authors have declared no competing interest.

## References

Agrawal, Puja, Steven Blinka, Kirthi Pulakanti, Michael H. Reimer, Cary Stelloh, Alison E. Meyer, and Sridhar Rao. 2021. “Genome Editing Demonstrates That the −5 Kb Nanog Enhancer Regulates Nanog Expression by Modulating RNAPII Initiation and/or Recruitment.” Journal of Biological Chemistry 296 (January): 100189. 10.1074/jbc.RA120.015152.

Alexander, Jeffrey M, Juan Guan, Bingkun Li, Lenka Maliskova, Michael Song, Yin Shen, Bo Huang, Stavros Lomvardas, and Orion D Weiner. 2019. “Live-Cell Imaging Reveals Enhancer-Dependent Sox2 Transcription in the Absence of Enhancer Proximity.” Edited by Robert H Singer, Kevin Struhl, and Zhe Liu. eLife 8 (May): e41769. 10.7554/eLife.41769.

Araki, K, M Araki, and K Yamamura. 1997. “Targeted Integration of DNA Using Mutant Lox Sites in Embryonic Stem Cells.” Nucleic Acids Research 25 (4): 868–72.

Arcot, S S, E K Flemington, and P L Deininger. 1989. “The Human Thymidine Kinase Gene Promoter.” Journal of Biological Chemistry 264 (4): 2343–49. 10.1016/S0021-9258(18)94182-7.

Arnold, Cosmas D., Daniel Gerlach, Christoph Stelzer, Łukasz M. Boryń, Martina Rath, and Alexander Stark. 2013. “Genome-Wide Quantitative Enhancer Activity Maps Identified by STARR-Seq.” *Science (New York*, N.Y.) 339 (6123): 1074–77. 10.1126/science.1232542.

Batut, Philippe J., Xin Yang Bing, Zachary Sisco, João Raimundo, Michal Levo, and Michael S. Levine. 2022. “Genome Organization Controls Transcriptional Dynamics during Development.” Science 375 (6580): 566–70. 10.1126/science.abi7178.

Benabdallah, Nezha S., Iain Williamson, Robert S. Illingworth, Lauren Kane, Shelagh Boyle, Dipta Sengupta, Graeme R. Grimes, Pierre Therizols, and Wendy A. Bickmore. 2019. “Decreased Enhancer-Promoter Proximity Accompanying Enhancer Activation.” Molecular Cell 76 (3): 473–484.e7. 10.1016/j.molcel.2019.07.038.

Blayney, Joseph, Helena Francis, Brendan Camellato, Leslie Mitchell, Rosa Stolper, Jef Boeke, Douglas Higgs, and Mira Kassouf. 2022. “Super-Enhancers Require a Combination of Classical Enhancers and Novel Facilitator Elements to Drive High Levels of Gene Expression.” bioRxiv. 10.1101/2022.06.20.496856.

Bothma, Jacques P., Hernan G. Garcia, Samuel Ng, Michael W. Perry, Thomas Gregor, and Michael Levine. 2015. “Enhancer Additivity and Non-Additivity Are Determined by Enhancer Strength in the Drosophila Embryo.” eLife 4. 10.7554/eLife.07956.

Branda, Catherine S., and Susan M. Dymecki. 2004. “Talking about a Revolution: The Impact of Site-Specific Recombinases on Genetic Analyses in Mice.” Developmental Cell 6 (1): 7–28. 10.1016/S1534-5807(03)00399-X.

Brosh, Ran, Camila Coelho, André M. Ribeiro-dos-Santos, Gwen Ellis, Megan S. Hogan, Hannah J. Ashe, Nicolette Somogyi, et al. 2023. “Synthetic Regulatory Genomics Uncovers Enhancer Context Dependence at the Sox2 Locus.” Molecular Cell, March, S1097276523001545. 10.1016/j.molcel.2023.02.027.

Buecker, Christa, Rajini Srinivasan, Zhixiang Wu, Eliezer Calo, Dario Acampora, Tiago Faial, Antonio Simeone, Minjia Tan, Tomasz Swigut, and Joanna Wysocka. 2014. “Reorganization of Enhancer Patterns in Transition from Naive to Primed Pluripotency.” Cell Stem Cell 14 (6): 838–53. 10.1016/j.stem.2014.04.003.

Cabrera, Alan, Hailey I. Edelstein, Fokion Glykofrydis, Kasey S. Love, Sebastian Palacios, Josh Tycko, Meng Zhang, et al. 2022. “The Sound of Silence: Transgene Silencing in Mammalian Cell Engineering.” Cell Systems 13 (12): 950–73. 10.1016/j.cels.2022.11.005.

Carleton, Julia B., Kristofer C. Berrett, and Jason Gertz. 2017. “Multiplex Enhancer Interference Reveals Collaborative Control of Gene Regulation by Estrogen Receptor α-Bound Enhancers.” Cell Systems 5 (4): 333–344.e5. 10.1016/j.cels.2017.08.011.

Catarino, Rui R., and Alexander Stark. 2018. “Assessing Sufficiency and Necessity of Enhancer Activities for Gene Expression and the Mechanisms of Transcription Activation.” Genes & Development 32 (3–4): 202–23. 10.1101/gad.310367.117.

Chen, Hongtao, Michal Levo, Lev Barinov, Miki Fujioka, James B. Jaynes, and Thomas Gregor. 2018. “Dynamic Interplay between Enhancer–Promoter Topology and Gene Activity.” Nature Genetics 50 (9): 1296–1303. 10.1038/s41588-018-0175-z.

Chen, Liang-Fu, Hannah Katherine Long, Minhee Park, Tomek Swigut, Alistair Nicol Boettiger, and Joanna Wysocka. 2023. “Structural Elements Promote Architectural Stripe Formation and Facilitate Ultra-Long-Range Gene Regulation at a Human Disease Locus.” *Molecular Cell*, March, S1097276523001661. 10.1016/j.molcel.2023.03.009.

Dixon, Jesse R., Siddarth Selvaraj, Feng Yue, Audrey Kim, Yan Li, Yin Shen, Ming Hu, Jun S. Liu, and Bing Ren. 2012. “Topological Domains in Mammalian Genomes Identified by Analysis of Chromatin Interactions.” Nature 485 (7398): 376–80. 10.1038/nature11082.

Gasperini, Molly, Jacob M. Tome, and Jay Shendure. 2020. “Towards a Comprehensive Catalogue of Validated and Target-Linked Human Enhancers.” Nature Reviews. Genetics 21 (5): 292–310. 10.1038/s41576-019-0209-0.

Gilpatrick, Timothy, Isac Lee, James E. Graham, Etienne Raimondeau, Rebecca Bowen, Andrew Heron, Bradley Downs, Saraswati Sukumar, Fritz J. Sedlazeck, and Winston Timp. 2020. “Targeted Nanopore Sequencing with Cas9-Guided Adapter Ligation.” Nature Biotechnology 38 (4): 433–38. 10.1038/s41587-020-0407-5.

Greenshpan, Yariv, Omri Sharabi, Ksenia M. Yegodayev, Ofra Novoplansky, Moshe Elkabets, Roi Gazit, and Angel Porgador. 2022. “The Contribution of the Minimal Promoter Element to the Activity of Synthetic Promoters Mediating CAR Expression in the Tumor Microenvironment.” International Journal of Molecular Sciences 23 (13): 7431. 10.3390/ijms23137431.

Gschwind, Andreas R., Kristy S. Mualim, Alireza Karbalayghareh, Maya U. Sheth, Kushal K. Dey, Evelyn Jagoda, Ramil N. Nurtdinov, et al. 2023. “An Encyclopedia of Enhancer-Gene Regulatory Interactions in the Human Genome.” bioRxiv. 10.1101/2023.11.09.563812.

Hay, Deborah, Jim R. Hughes, Christian Babbs, James O. J. Davies, Bryony J. Graham, Lars L. P. Hanssen, Mira T. Kassouf, et al. 2016. “Genetic Dissection of the α-Globin Super-Enhancer in Vivo.” Nature Genetics 48 (8): 895–903. 10.1038/ng.3605.

Heist, Tyler, Takashi Fukaya, and Michael Levine. 2019. “Large Distances Separate Coregulated Genes in Living Drosophila Embryos.” Proceedings of the National Academy of Sciences 116 (30): 15062–67. 10.1073/pnas.1908962116.

Hnisz, Denes, Jurian Schuijers, Charles Y. Lin, Abraham S. Weintraub, Brian J. Abraham, Tong Ihn Lee, James E. Bradner, and Richard A. Young. 2015. “Convergence of Developmental and Oncogenic Signaling Pathways at Transcriptional Super-Enhancers.” Molecular Cell 58 (2): 362–70. 10.1016/j.molcel.2015.02.014.

Huang, Jialiang, Xin Liu, Dan Li, Zhen Shao, Hui Cao, Yuannyu Zhang, Eirini Trompouki, et al. 2016. “Dynamic Control of Enhancer Repertoires Drives Lineage and Stage-Specific Transcription during Hematopoiesis.” Developmental Cell 36 (1): 9–23. 10.1016/j.devcel.2015.12.014.

Kim, Seungsoo, and Joanna Wysocka. 2023. “Deciphering the Multi-Scale, Quantitative Cis-Regulatory Code.” *Molecular Cell*, Reimagining the Central Dogma, 83 (3): 373–92. 10.1016/j.molcel.2022.12.032.

Li, Jieru, Ankun Dong, Kamola Saydaminova, Hill Chang, Guanshi Wang, Hiroshi Ochiai, Takashi Yamamoto, and Alexandros Pertsinidis. 2019. “Single-Molecule Nanoscopy Elucidates RNA Polymerase II Transcription at Single Genes in Live Cells.” Cell 178 (2): 491–506.e28. 10.1016/j.cell.2019.05.029.

Li, Jieru, Angela Hsu, Yujing Hua, Guanshi Wang, Lingling Cheng, Hiroshi Ochiai, Takashi Yamamoto, and Alexandros Pertsinidis. 2020. “Single-Gene Imaging Links Genome Topology, Promoter-Enhancer Communication and Transcription Control.” Nature Structural & Molecular Biology 27 (11): 1032–40. 10.1038/s41594-020-0493-6.

Lienert, Florian, Christiane Wirbelauer, Indrani Som, Ann Dean, Fabio Mohn, and Dirk Schübeler. 2011. “Identification of Genetic Elements That Autonomously Determine DNA Methylation States.” Nature Genetics 43 (11): 1091–97. 10.1038/ng.946.

Long, Hannah K., Sara L. Prescott, and Joanna Wysocka. 2016. “Ever-Changing Landscapes: Transcriptional Enhancers in Development and Evolution.” Cell 167 (5): 1170–87. 10.1016/j.cell.2016.09.018.

Loubiere, Vincent, Bernardo P. de Almeida, Michaela Pagani, and Alexander Stark. 2023. “Developmental and Housekeeping Transcriptional Programs Display Distinct Modes of Enhancer-Enhancer Cooperativity in Drosophila.” bioRxiv. 10.1101/2023.10.10.561770.

Martinez-Ara, Miguel, Federico Comoglio, and Bas van Steensel. 2023. “Large-Scale Analysis of the Integration of Enhancer-Enhancer Signals by Promoters.” bioRxiv. 10.1101/2023.08.11.552995.

Moorthy, Sakthi D., Scott Davidson, Virlana M. Shchuka, Gurdeep Singh, Nakisa Malek-Gilani, Lida Langroudi, Alexandre Martchenko, Vincent So, Neil N. Macpherson, and Jennifer A. Mitchell. 2017. “Enhancers and Super-Enhancers Have an Equivalent Regulatory Role in Embryonic Stem Cells through Regulation of Single or Multiple Genes.” Genome Research 27 (2): 246–58. 10.1101/gr.210930.116.

Muerdter, Felix, Łukasz M. Boryń, Ashley R. Woodfin, Christoph Neumayr, Martina Rath, Muhammad A. Zabidi, Michaela Pagani, et al. 2018. “Resolving Systematic Errors in Widely Used Enhancer Activity Assays in Human Cells.” Nature Methods 15 (2): 141–49. 10.1038/nmeth.4534.

Neumayr, Christoph, Vanja Haberle, Leonid Serebreni, Katharina Karner, Oliver Hendy, Ann Boija, Jonathan E. Henninger, et al. 2022. “Differential Cofactor Dependencies Define Distinct Types of Human Enhancers.” Nature 606 (7913): 406–13. 10.1038/s41586-022-04779-x.

Olbrich, Teresa, Maria Vega-Sendino, Desiree Tillo, Wei Wu, Nicholas Zolnerowich, Raphael Pavani, Andy D. Tran, et al. 2021. “CTCF Is a Barrier for 2C-like Reprogramming.” Nature Communications 12 (1): 4856. 10.1038/s41467-021-25072-x.

Pachano, Tomás, Endika Haro, and Alvaro Rada-Iglesias. 2022. “Enhancer-Gene Specificity in Development and Disease.” Development 149 (11): dev186536. 10.1242/dev.186536.

Patty, Benjamin J., and Sarah J. Hainer. 2021. “Transcription Factor Chromatin Profiling Genome-Wide Using uliCUT&RUN in Single Cells and Individual Blastocysts.” Nature Protocols 16 (5): 2633–66. 10.1038/s41596-021-00516-2.

Peng, Tianran, Yanan Zhai, Yaser Atlasi, Menno ter Huurne, Hendrik Marks, Hendrik G. Stunnenberg, and Wout Megchelenbrink. 2020. “STARR-Seq Identifies Active, Chromatin-Masked, and Dormant Enhancers in Pluripotent Mouse Embryonic Stem Cells.” Genome Biology 21 (1): 243. 10.1186/s13059-020-02156-3.

Rao, Suhas S. P., Miriam H. Huntley, Neva C. Durand, Elena K. Stamenova, Ivan D. Bochkov, James T. Robinson, Adrian L. Sanborn, et al. 2014. “A 3D Map of the Human Genome at Kilobase Resolution Reveals Principles of Chromatin Looping.” Cell 159 (7): 1665–80. 10.1016/j.cell.2014.11.021.

Rideout, William M., Teruhiko Wakayama, Anton Wutz, Kevin Eggan, Laurie Jackson-Grusby, Jessica Dausman, Ryuzo Yanagimachi, and Rudolf Jaenisch. 2000. “Generation of Mice from Wild-Type and Targeted ES Cells by Nuclear Cloning.” Nature Genetics 24 (2): 109–10. 10.1038/72753.

Rinzema, Niels J., Konstantinos Sofiadis, Sjoerd J. D. Tjalsma, Marjon J. A. M. Verstegen, Yuva Oz, Christian Valdes-Quezada, Anna-Karina Felder, et al. 2022. “Building Regulatory Landscapes Reveals That an Enhancer Can Recruit Cohesin to Create Contact Domains, Engage CTCF Sites and Activate Distant Genes.” Nature Structural & Molecular Biology 29 (6): 563–74. 10.1038/s41594-022-00787-7.

Shin, Ha Youn, Michaela Willi, Kyung Hyun Yoo, Xianke Zeng, Chaochen Wang, Gil Metser, and Lothar Hennighausen. 2016. “Hierarchy within the Mammary STAT5-Driven Wap Super-Enhancer.” Nature Genetics 48 (8): 904–11. 10.1038/ng.3606.

Smith, Robin P., Leila Taher, Rupali P. Patwardhan, Mee J. Kim, Fumitaka Inoue, Jay Shendure, Ivan Ovcharenko, and Nadav Ahituv. 2013. “Massively Parallel Decoding of Mammalian Regulatory Sequences Supports a Flexible Organizational Model.” Nature Genetics 45 (9): 1021–28. 10.1038/ng.2713.

Taylor, Tiegh, Natalia Sikorska, Virlana M. Shchuka, Sanjay Chahar, Chenfan Ji, Neil N. Macpherson, Sakthi D. Moorthy, et al. 2022. “Transcriptional Regulation and Chromatin Architecture Maintenance Are Decoupled Functions at the Sox2 Locus.” Genes & Development 36 (11–12): 699–717. 10.1101/gad.349489.122.

Thomas, Henry F., Elena Kotova, Swathi Jayaram, Axel Pilz, Merrit Romeike, Andreas Lackner, Thomas Penz, et al. 2021. “Temporal Dissection of an Enhancer Cluster Reveals Distinct Temporal and Functional Contributions of Individual Elements.” Molecular Cell 81 (5): 969–982.e13. 10.1016/j.molcel.2020.12.047.

Weinberg, Daniel N., Simon Papillon-Cavanagh, Haifen Chen, Yuan Yue, Xiao Chen, Kartik N. Rajagopalan, Cynthia Horth, et al. 2019. “The Histone Mark H3K36me2 Recruits DNMT3A and Shapes the Intergenic DNA Methylation Landscape.” Nature 573 (7773): 281–86. 10.1038/s41586-019-1534-3.

Wit, Elzo de, Erica S. M. Vos, Sjoerd J. B. Holwerda, Christian Valdes-Quezada, Marjon J. A. M. Verstegen, Hans Teunissen, Erik Splinter, Patrick J. Wijchers, Peter H. L. Krijger, and Wouter de Laat. 2015. “CTCF Binding Polarity Determines Chromatin Looping.” Molecular Cell 60 (4): 676–84. 10.1016/j.molcel.2015.09.023.

Yang, Pengyi, Sean J. Humphrey, Senthilkumar Cinghu, Rajneesh Pathania, Andrew J. Oldfield, Dhirendra Kumar, Dinuka Perera, et al. 2019. “Multi-Omic Profiling Reveals Dynamics of the Phased Progression of Pluripotency.” Cell Systems 8 (5): 427–445.e10. 10.1016/j.cels.2019.03.012.

Zabidi, Muhammad A., Cosmas D. Arnold, Katharina Schernhuber, Michaela Pagani, Martina Rath, Olga Frank, and Alexander Stark. 2015. “Enhancer-Core-Promoter Specificity Separates Developmental and Housekeeping Gene Regulation.” Nature 518 (7540): 556–59. 10.1038/nature13994.

Zuin, Jessica, Gregory Roth, Yinxiu Zhan, Julie Cramard, Josef Redolfi, Ewa Piskadlo, Pia Mach, et al. 2022. “Nonlinear Control of Transcription through Enhancer–Promoter Interactions.” Nature 604 (7906): 571–77. 10.1038/s41586-022-04570-y.

